# Decoding Dynamically Shifting States of Parkinson’s Disease: Tremor, Bradykinesia, and Effective Motor Control

**DOI:** 10.1101/2022.02.12.480213

**Authors:** Peter M. Lauro, Shane Lee, Daniel E. Amaya, David D. Liu, Umer Akbar, Wael F Asaad

## Abstract

Parkinson’s Disease (PD) is characterized by distinct motor phenomena that are expressed asynchronously. Understanding the neurophysiological correlates of these different motor states could facilitate monitoring of disease progression and allow improved assessments of therapeutic efficacy, as well as enable optimal closed-loop neuromodulation. We examined neural activity in the basal ganglia and cortex of subjects with PD during a quantitative motor task to decode tremor and bradykinesia — two cardinal motor signs of this disease — and relatively asymptomatic periods of behavior. Analysis of subcortical and cortical signals revealed that tremor and bradykinesia had distinct, nearly opposite neural signatures, while effective motor control displayed unique, differentiating features. The neurophysiological signatures of these motor states depended on the type of signal recorded as well as the location; cortical decoding accuracy generally outperformed subcortical decoding, while tremor and bradykinesia were better decoded from different portions of the subthalamic nucleus (STN). These results provide a roadmap to leverage real-time neurophysiology to understand and treat PD.

**One Sentence Summary:** Motor signs of Parkinson’s disease such as tremor and bradykinesia can be independently decoded from subthalamic and cortical recordings.

## INTRODUCTION

Parkinson’s disease (PD) is a common and complex neurodegenerative disorder characterized by the dynamic expression of particular motor features such as tremor and bradykinesia *(1, 2)*. These distinct motor signs are expressed variably across patients and may respond differently to dopamine replacement therapy; their differential expression often is used to classify patients into phenotypic subtypes *(3, 4)*. Despite this heterogeneity, both of these motor features (and both tremor-dominant and non-tremor-dominant patient subtypes) respond to high-frequency deep brain stimulation (DBS) applied to the subthalamic nucleus (STN) *(5, 6)*.

DBS delivered in a closed-loop fashion (*i*.*e*., in response to neurophysiological biomarkers) has shown promising therapeutic potential primarily toward alleviating bradykinesia *(7, 8)*, but current efforts focusing on *β*-frequency oscillations (15–30 Hz) have been shown rather to worsen tremor in many cases *(9)*. Thus, tremor may be better signaled by different components within the local field potential (LFP) spectrum, and closed-loop DBS strategies could benefit from a clearer understanding of the neurophysiological biomarkers that differentiate these motor signs from each other, and from more optimal motor performance in the absence of these impairments.

To this point, STN LFP recordings from patients with different PD subtypes have revealed distinct patterns of oscillatory activity *(10)*. In addition to spectral variability, specific stimulation sites within the STN have been associated with the preferential reduction of individual motor signs *(11)*. Moreover, these STN sites were associated with specific patterns of anatomical connectivity with cortical structures *(12)*. Much like how overlapping subdivisions of basal-ganglia-cortical circuits have been found to encode separate aspects of movement *(13, 14)*, separate motor features may be mediated by different sub-circuits involving the STN and sensorimotor cortex *(15)*.

In order to better reveal the functional and anatomical substrates of distinct PD motor states, we enlisted patients with PD undergoing awake DBS electrode implantation to perform a continuous visual-motor task that allowed rigorous, concurrent measurement of different motor metrics while we acquired STN (micro- and macro-electrode) and cortical (electrocorticography; ECoG) recordings. Prior studies have not attempted to simultaneously decode different aspects of disease expression, contrast these measures with symptom-free performance, and examine disease expression on the short timescales relevant to that varying expression. While our group has previously demonstrated the ability to decode global PD motor dysfunction from STN recordings on short timescales *(16, 17)*, we focus here on individual motor features and their specific neurophysiological manifestations. Specifically, we trained machine learning models to directly decode tremor or slowness from neural recordings to reveal the spectral and anatomical fingerprints of these cardinal motor features of PD.

## RESULTS

### Motor behavior during the target tracking task

Twenty-seven patients with Parkinson’s disease (PD) undergoing STN DBS implantation and 17 age-matched controls (see *Materials and Methods, Study and Experimental Design and Behavior Metric Analysis*) performed a visual-motor task in which they followed an on-screen target with a cursor controlled by either a joystick or a stylus and tablet **(Fig. 1A)**. Twenty-three patients (and 12 control subjects) performed a version of the task with fixed-patterns of target movement, while 4 patients (and 5 control subjects) performed a version with randomly-generated target paths unique to each trial. Each patient performed 1–4 sessions of this target-tracking task during the procedure for a total of 69 sessions, while control subjects each performed 1 session extra-operatively for a total of 17 sessions. Tremor amplitude and cursor speed, task metrics calculated to reflect the expression of tremor and bradykinesia, were quantified continuously from the x- and y-cursor traces. These behavioral metric data were then averaged into 7 second non-overlapping epochs. To compare metrics across subject populations while controlling for the unequal epoch contributions of each subject, linear mixed models (LMMs) were used (see *Materials and Methods, Statistical Analysis*). The resulting distributions for PD vs. control subjects across task versions demonstrated increased tremor for PD patients (*n* = 6498 epochs across 44 subjects, PD vs. Control, linear mixed model coefficient/LMM *β* = 0.337, *Z* = 2.169, *p* = 0.030), but only a trend for decreased speed (PD vs. Control, LMM *β* = −0.592, *Z* = −1.194, *p* = 0.232) **(Fig. 1B, C)**.

**Fig. 1.**
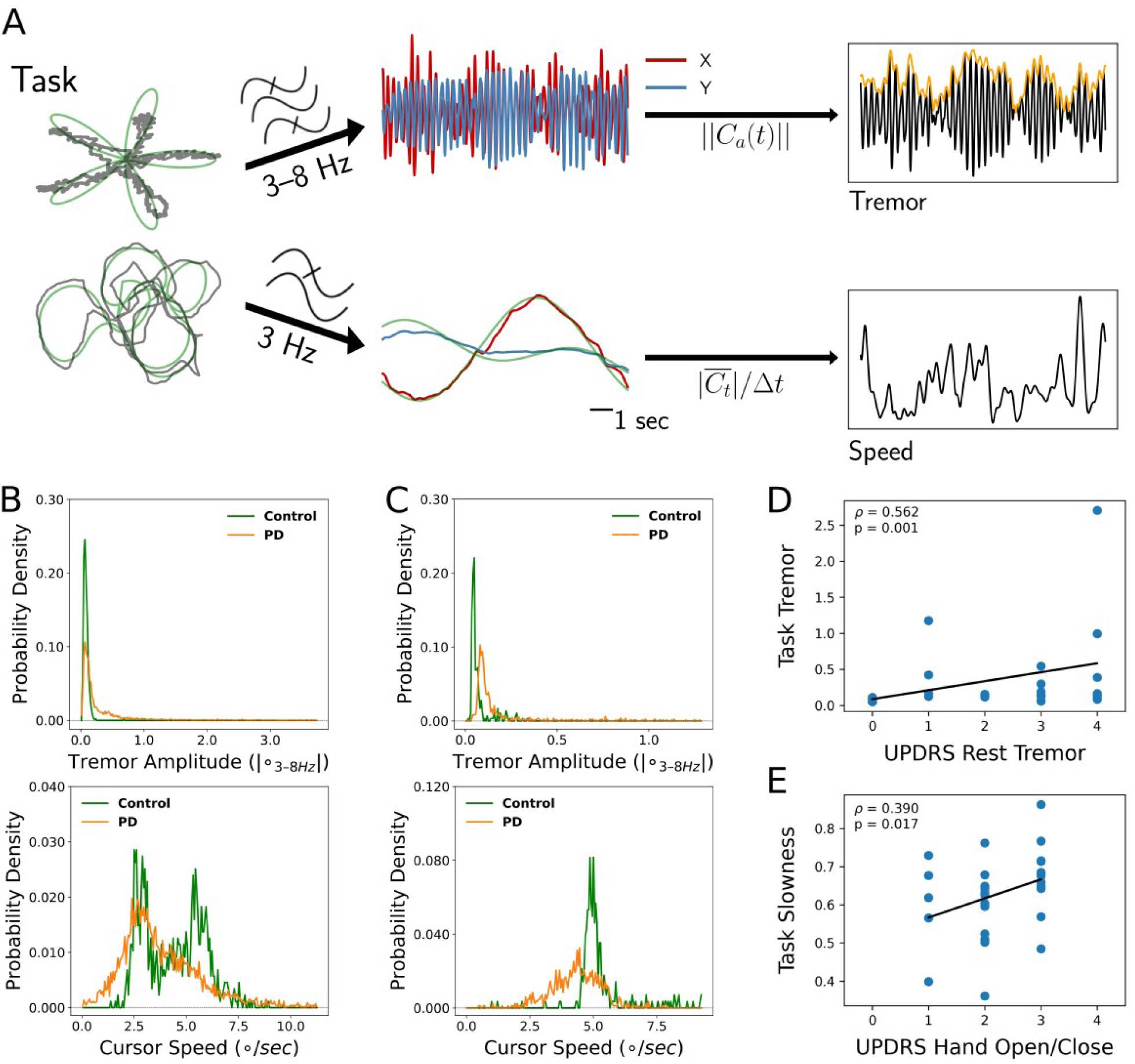
Tremor and movement speed calculated from fixed- and random-pattern intraoperative visual-motor tasks. **(A)** Left-top: Schematic of task target (green) and cursor (gray) traces from a single trial of the fixed-pattern task. Left-bottom: Schematic of task target (green) and cursor (gray) traces from a single trial of the random-pattern task. Center-top: Bandpass filtered X and Y cursor traces from a task trial. ∥*C*_*a*_∥ refers to the amplitude of the analytic signal (*a*) of the cursor trace (*C*). Center-bottom: Lowpass filtered X and Y cursor traces from a task trial. Right-top: One-dimensional projection of bandpass filtered traces (black), with tremor amplitude measured from the envelope (orange). Right-bottom: Cursor speed measured from lowpass filtered traces (black). Figure adapted from *(17)*. **(B)** Distributions of 7 second tremor amplitude (top) and cursor speed (bottom) epochs for control subject and PD patient populations in the fixed-pattern task (*n* = 23 patients with PD, *n* = 12 age-matched control subjects). ° - degrees of visual angle. While there was overlap on the left side of the tremor distribution (patients with PD can exhibit control-like levels of tremor), the PD tremor distribution was highly skewed on the right side of the distribution, demonstrating a large range of tremor expression. The bimodality of the control speed distribution corresponded to the pre-programmed speed of the onscreen target. Despite this, note that the PD speed distribution was shifted towards lower speed values. **(C)** Distributions of 7 second tremor amplitude (top) and cursor speed (bottom) epochs for control subject and PD patient populations in the random-pattern task (*n* = 4 patients with PD, *n* = 5 age-matched control subjects). Note that the control population speed was unimodal, corresponding to the fixed speed of the onscreen target across all random patterns. Again, note that the PD speed distribution was shifted towards lower speed values. **(D)** Tremor amplitude (as measured by the tasks) corresponded to UPDRS measures of tremor. *ρ* = Spearman’s correlation statistic and its associated p value. **(E)** Movement slowness (as measured by the tasks) corresponded to UPDRS measures of bradykinesia.

Qualitatively, the distributions of tremor magnitude in these populations were similar regardless of task versions, indicating that subjects with PD spent a substantial fraction of time in states resembling controls, but the PD the distribution demonstrated a long tail, indicating a large range of tremor expression in patients not present in control subjects in both task versions **(Fig. 1B, C; top)**. On the other hand, the two task versions generated different movement speed distributions **(Fig. 1B, C; bottom)**. While the fixed-pattern version of the task elicited a bimodal distribution (reflecting slower turns and faster straight path segments), the random-pattern version elicited a single peak corresponding to the fixed target speed used in that task. Nonetheless, in both versions, the PD cursor speed distribution was shifted to the left (*i*.*e*., slower) relative to control subjects. Cursor speed was subsequently converted to “slowness” by normalizing each session’s distribution of cursor speed to its minimum and maximum values (0: highest speed, 1: lowest speed). To determine whether these two motor metrics reflected distinct components of PD motor dysfunction, each patient’s metric distribution medians were correlated against their UPDRS III motor sub-scores. Indeed, tremor amplitude positively correlated with both the resting and postural tremor subscores (*ρ* =0.562, *p*=0.028, *n* = 27 PD subjects, Spearman correlation) **(Fig. 1D)** and slowness positively correlated with the hand open/close subscore (a subscore commonly used in part to assess bradykinesia) (*ρ* =0.390, *p*=0.014, *n* = 27 PD subjects, Spearman correlation) **(Fig. 1E)**. Thus, these two metrics reflected two specific aspects of Parkinson’s disease motor dysfunction.

### Tremor and slowness were distinct and opposing behavioral states

Relative to each other, tremor and slowness typically did not co-occur but rather were inversely expressed in time (tremor vs. slowness, LMM *β* = −0.552 across all sessions, *Z* = −20.551, *p* = 7.479 * 10^−94^) **(Fig. 2A, B)**. To understand whether this observed anti-correlation may have been due in part to motor features manifesting on different time scales, the autocorrelogram was computed for each metric with 100 ms epochs. Here, we found that (as determined by the autocorrelogram full-width half-maximum) tremor was typically expressed continuously for longer periods (0.898 s) as opposed to speed (0.297 s) **(Fig. 2C)**. Using this FWHM as the minimum, we calculated periods of time where metrics were sustained above control levels (*i*.*e*., symptomatic periods) across subjects. We found that the median duration of symptomatic tremor episodes was 2.000 s, and slowness episodes lasted for 0.500 s **(Fig. 2D)**. With these differing timescales and anti-correlated presence, we understood tremor and slowness to represent distinct behavioral states.

**Fig. 2.**
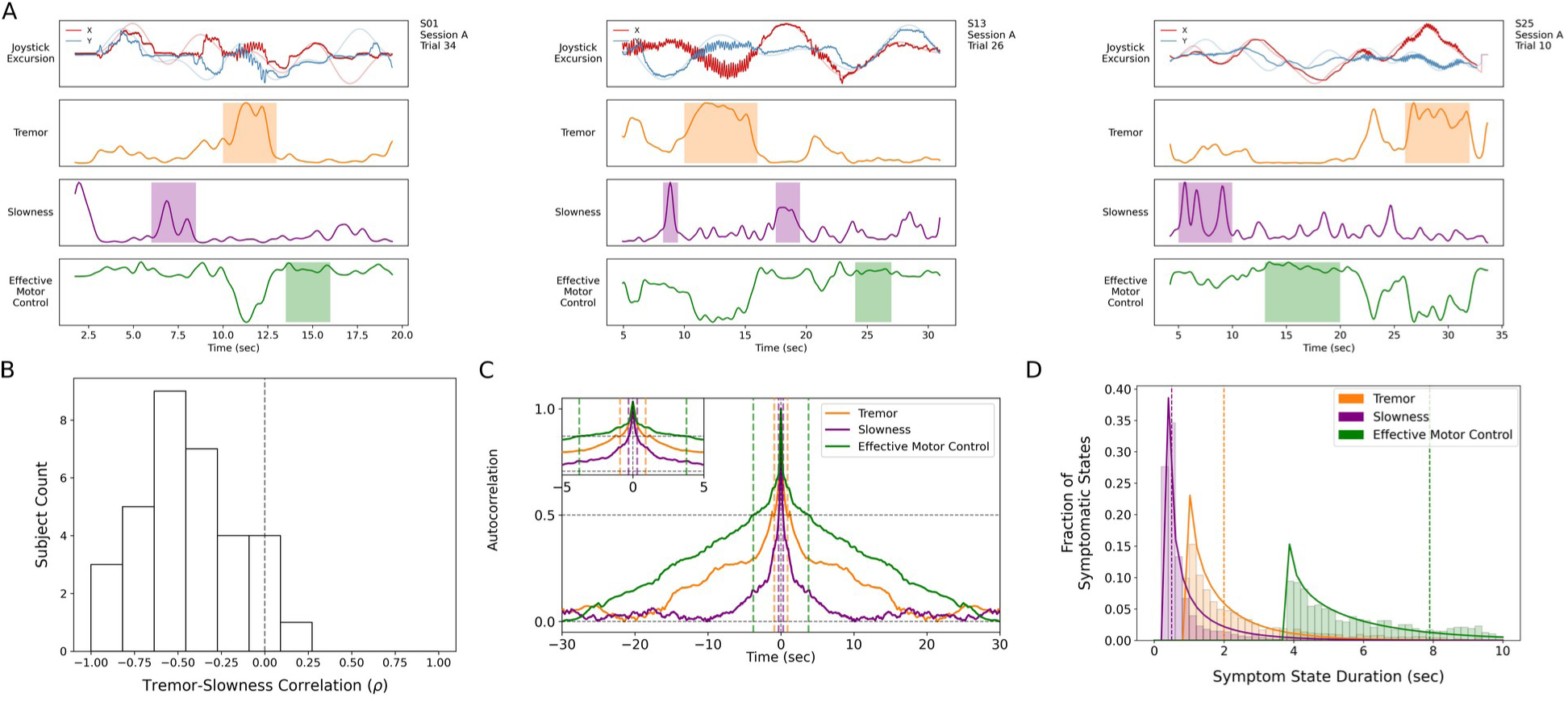
Tremor and slowness represented two non-overlapping motor states with differing timescales. **(A)** Examples from three individual subjects of behavioral (solid lines) and target (translucent lines) traces (top row) and calculated motor metrics (bottom three rows) within single trials. Periods of increased expression of individual motor metrics are highlighted by their respective color within the timeseries. Note how individual motor metrics often do not co-express. **(B)** Histogram of subject-wide behavioral Spearman correlation with tremor and slowness metrics. Vertical dashed line indicates 0. **(C)** Autocorrelograms of symptomatic (tremor, slowness) and non-symptomatic (effective motor control) metrics. Colored vertical dashed lines indicate full-width half-maximum (FWHM) for each metric. Top-left inset depicts a zoomed-in window of the autocorrelogram. **(D)** Histogram of sustained motor metric period duration (*i*.*e*., symptomatic state duration) across subjects. Solid lines indicate gamma distribution fit to each motor metric state histogram, while dashed vertical lines indicate the median symptomatic state length for each metric.

### Tremor and slowness had distinct representations within the subthalamic nucleus

A total of 203 microelectrode and 176 macroelectrode recordings were acquired from the region of the STN as patients performed the task. To directly assess whether tremor or slowness could be decoded from these deep recordings (and if one metric could be better decoded than the other), spectral estimates of power from 3–400 Hz were obtained using a wavelet convolution. Narrowband power estimates were grouped into six broad frequency bands (*θ* / *α, β, γ* _*low*_, *γ*_*mid*_, *γ*_*high*_, hfo) with 7 sub-bands each, for a total of 42 neural “features” per 7-second epoch *(17)*. Neural decoding models (support vector regression/SVR with a linear kernel) were trained directly on the epoch’s average metric (tremor or slowness values averaged within each epoch), and their performance was assessed with Pearson’s *r* between observed and decoded metrics. Across all microelectrode recordings (that is, regardless of precise recording location within or near the STN), tremor decoding performance (*r*=0.196 ± 0.250) was superior to slowness decoding (*r*=0.114 ± 0.193) (*n* = 203 microelectrode recordings, tremor v. slowness, LMM *β* = 0.083, *Z* = 4.626, *p* = 4.000*10^−6^). However, no such difference was observed across macroelectrode recordings (tremor v. slowness, *r*=0.208 ±0.283 v. *r*=0.217 ± 0.274, *n* = 176 macroelectrode recordings, LMM *β* = −0.009, *Z* = −0.557, *p* = 0.578).

To determine if tremor and slowness had distinct neurophysiological signatures, SVR model feature weights were computed for individual models and aggregated for each metric. First, to understand which spectral features were used consistently for individual metric models, feature weights were compared to null distributions generated from separate models in which motor metric values were shuffled with respect to the corresponding spectral features. Microelectrode tremor decoding models (*n* = 203 microelectrode recording models) consistently and positively weighted low-frequency features (*θ, α, β*; 4–21 Hz; *p*<0.001, permutation test, see *Methods*) **(Fig. 3A, left)**. To a lesser extent, *hfo* (275–375 Hz) weights were also positively associated with tremor decoding (p < 0.014, permutation test). In contrast, macroelectrode tremor decoding models (*n* = 176 macroelectrode recording models) negatively weighed *β* power (14–41 Hz) (p < 0.026, permutation test) while more clearly positively weighing broadband *γ*/*hfo* activity (60– 375 Hz) (*p*<0.011, permutation test) **(Fig. 3A, right)**. In other words, optimal macroelectrode tremor decoding relied on decreased *β* power and increased *γ*/*hfo* power.

**Fig. 3.**
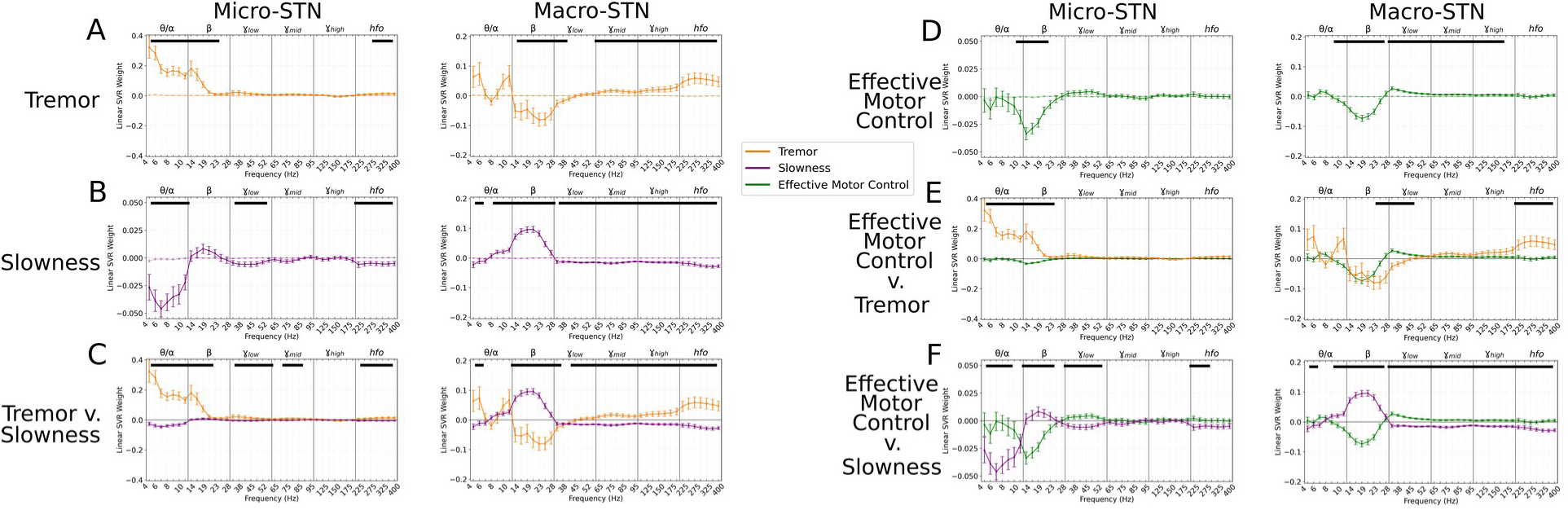
Subthalamic tremor decoding models emphasized lower frequencies whereas slowness models emphasized higher frequencies. **(A)** Average tremor decoding model coefficients for every STN microelectrode (left) and macroelectrode (right) recording. Solid colored lines indicate average weights, with positive/negative values reflecting a positive or negative relationship with tremor amplitude. Error bars indicate s.e.m. across subjects. Translucent colored lines represented mean weights (± s.e.m.) for SVR model weights calculated on shuffled metric data. Black solid lines (top) represented contiguous spectral features that were observed above chance. **(B)** Average slowness decoding model coefficients for every STN microelectrode (left) and macroelectrode (right) recording. **(C)** Average model coefficients for tremor and slowness for all STN microelectrode (left) and macroelectrode (right) recordings. Black solid lines (top) represented contiguous spectral features that significantly differed between tremor and slowness decoding models. **(D)** Average effective motor control decoding model coefficients for every STN microelectrode (left) and macroelectrode (right) recording. **(E)** Average model coefficients for effective motor control and tremor for all STN microelectrode (left) and macroelectrode (right) recordings. **(F)** Average model coefficients for effective motor control and slowness for all STN microelectrode (left) and macroelectrode (right) recordings.

For slowness, microelectrode decoding models consistently had negative *θ, γ*_*low*_, and *hfo* weights (5–12 Hz; 33–56 Hz; 200–375 Hz) (*p*<0.012, permutation test) **(Fig. 3B, left)**. While *β* did not significantly modulate with slowness in microelectrode decoding models, positive *β* (12–30 Hz) weights were observed in macroelectrode decoding models (p < 0.006, permutation test), along with negative *γ*/*hfo* weights (33–375 Hz) (p < 0.001, permutation test) **(Fig. 3B, right)**. Tremor and slowness model features were also found to differ from each other when compared directly, with *hfo* frequencies (225–375 Hz) being elevated during tremor in both micro-/macro-electrode recordings **(Fig. 3C)**. Overall, just as tremor and slowness represented two distinct, anti-correlated behavioral states of Parkinson’s disease motor dysfunction, tremor and slowness decoding models from the STN revealed clearly distinguishable patterns of underlying neural activity.

However, in order to rule out the possibility that the alternating patterns of relevant neural decoding features simply reflected the anti-correlated nature of tremor and slowness, we tested whether decoding models trained for tremor could accurately decode slowness. When directly comparing the performance of tremor and slowness decoding on tremor-trained models, slowness decoding was inferior for both microelectrode (tremor v. slowness decoding, *r* = 0.194 ± 0.253 v. 0.000 ± 0.025; LMM β = 0.195, *Z* = 12.373 *p* = 3.67*10^−35^) and macroelectrode (tremor v. slowness decoding, *r* = 0.205 ± 0.213 v. −0.005 ± 0.024; LMM β = 0.210, *Z* = 14.360, *p* = 9.28*10^−47^) recordings. If decoding features for tremor and slowness were simply inverted, applying models for decoding another metric would result in negative *r*-values above chance. Thus, the neural features used for individual metric decoding likely reflected a unique spectral state or “fingerprint,” not simply the correlated presence or absence of motor features.

### Effective motor control had characteristic neural signatures

PD produces a fluctuating motor deficit such that there can be moments of normal-appearing motor behavior *(18)*. We therefore asked whether there existed a distinguishable pattern of neural activity that represented these “effective” motor states. Specifically, epochs with lower tremor and/or higher movement speeds were assigned values closer to 1, while more symptomatic epochs (high tremor and/or slower movement speeds) were assigned values closer to 0. When comparing the timescale of effective motor control to other metrics, we found it to be expressed on longer timescales (FWHM = 3.784 s, median state length = 7.900 s) **(Fig. 2A, C, D)**.

Applying the same decoding methods as above, we found that effective motor control could be decoded above chance from both micro- (*r* = 0.138 ± 0.194) and macro-electrode (*r* = 0.228 ± 0.181) recordings. Further, effective motor control was found to be characterized by the absence of *β* (10–28 Hz) power in both micro- and macro-electrode recordings (*p*<0.006, permutation test), while macroelectrodes also exhibited positive *γ* power weights (30–175 Hz) (*p*<0.020, permutation test) **(Fig. 3D)**. Power in *γ*_*low*_ frequencies (30–48 Hz) in particular was found to be significantly increased during effective motor control relative to both tremor and slowness decoding models (*p*<0.006, permutation test) **(Fig. 3E-F, right)**.

In total, STN activity contained specific features that distinguished symptomatic from non-symptomatic motor states. Tremor was characterized by lower frequencies (*θ* / *α*) in microelectrodes, slowness by *β* frequencies in macroelectrodes, and effective motor performance was uniquely characterized by *γ*_*low*_ frequencies from both recording types.

### Optimal subthalamic tremor decoding sites were dorsolateral to optimal slowness-decoding sites across patients

To investigate whether tremor and slowness were more optimally decoded from distinct areas within the STN, recording sites for each session were reconstructed using subject-specific neuroimaging (peak microelectrode recording density in MNI space: *x*=−12, *y*=−10, *z*=−6.0) **(Fig. 4A)**; peak macroelectrode recording density in MNI space: *x*=−12, *y*=−9, *z*=−3.0). For each recording site, the corresponding decoding model performance for each metric was plotted, and the differences between motor feature SVR *r*-values were then calculated per voxel.

**Fig. 4.**
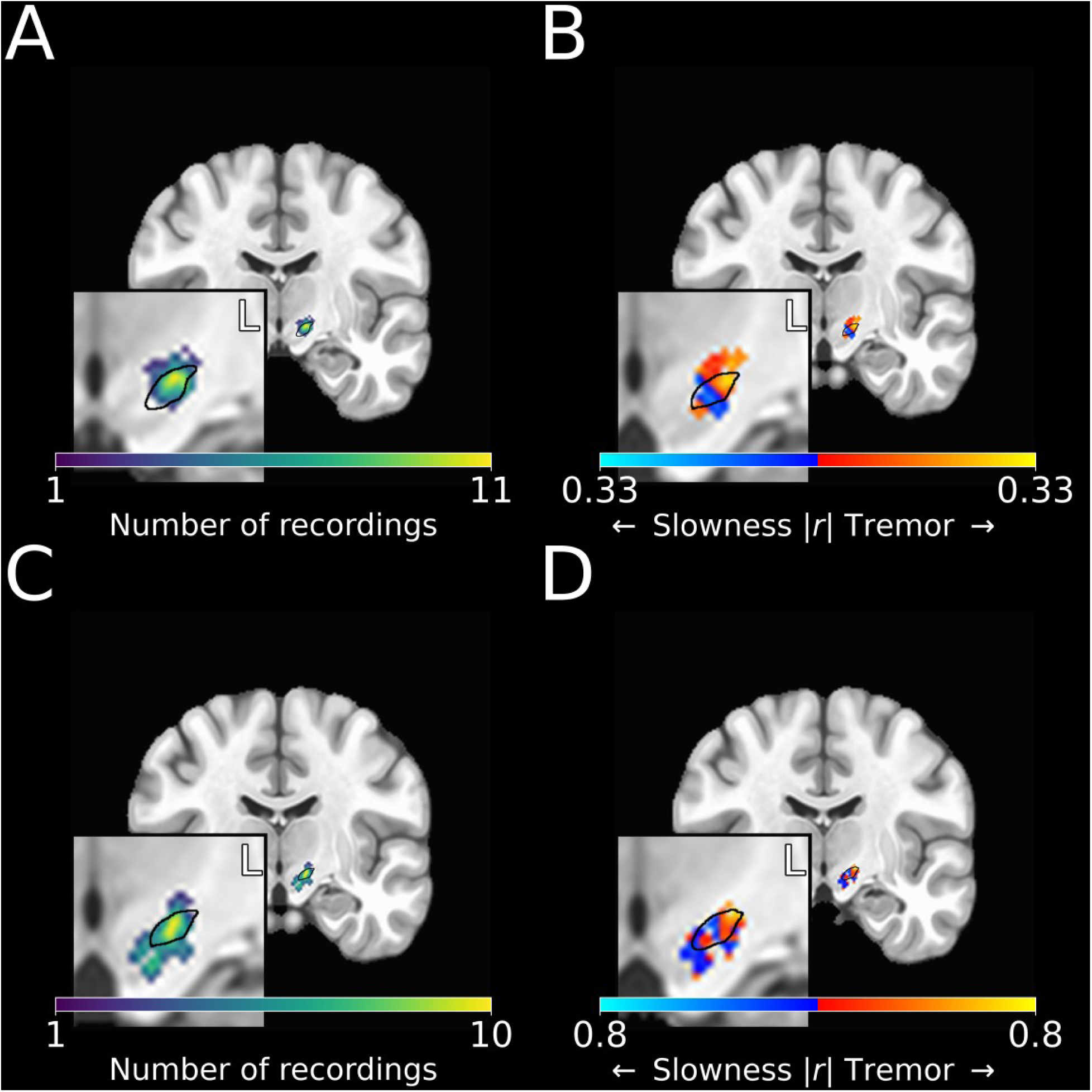
Optimal subthalamic tremor-decoding sites were dorsolateral to optimal slowness-decoding sites. **(A)** Recording density of stationary microelectrode recordings across all patients and task sessions overlaid on an MNI reference volume (approximate outline of the STN in black). L: left. **(B)** Difference in tremor vs. slowness decoding *r*-values for stationary microelectrode recordings. Warmer colors indicate voxels where tremor decoding was superior, whereas cooler colors indicate where slowness decoding was superior. Note that tremor was better decoded in more dorsolateral STN, with better slowness decoding observed in central/ventromedial STN. **(C)** Recording density of moving microelectrode recordings across all patients and task sessions overlaid on an MNI reference volume (approximate outline of the STN in black). L: left. **(D)** Difference in tremor vs. slowness decoding *r*-values for high-density STN survey recordings. *r-*values depicted here are site-specific *r-*values generated from the whole-STN model applied to individual depth recordings. Warmer colors indicate voxels where tremor decoding was superior, whereas cooler colors indicate where slowness decoding was superior. Despite the results of the high-density STN survey recording results (*n* = 5 patients) being noisier than those of stationary microelectrode recording results (*n* = 27 patients), tremor and slowness were optimally decoded in similar areas of the STN.

Across microelectrode recordings, tremor was optimally decoded at *x*=−14.0, *y*=−13.0, *z*=−5.0 **(Fig. 4B)**. Slowness on the other hand was optimally decoded at *x*=−11.0, *y*=−13.0, *z*=−8.0. We then directly compared the voxel-wise relative performance between tremor and slowness throughout all recorded STN voxels by using a modified 3D *t*-test which derived voxel-wise *Z*-statistics and *p*-values from 10000 permutations of randomly shuffling *r*-values among voxels (“3dttest++” with “-ETAC” option, see *Methods*). Using this, we found that tremor was better decoded in recordings from dorsolateral STN, whereas slowness was better decoded from recordings in central/ventromedial STN. Both tremor (*n* = 203 microelectrode recording sites, AFNI’s “3dttest++”, *Z*=2.207, *p*=0.014) and slowness (*Z*=1.915, *p*=0.0 28) decoding peaks were observed above chance.

However, optimal locations for tremor and slowness decoding were not found to differ significantly when assessed using the broader spatial sampling afforded by macroelectrode location (*n* = 176 macroelectrode recording sites, AFNI’s “3dttest++”, *p*>0 .05). Moreover, the locus of optimal effective motor control decoding was not observed to differ from those of tremor or speed using either micro- and macro-electrode recordings (AFNI’s “3dttest++”, *p*>0.05). Nevertheless, based on the higher anatomical precision of microelectrode recordings, we found that that differences in metric decoding were not only related to the frequencies present, but also to an electrode’s precise location within the STN, as assessed over the entire study PD population.

### Optimal subthalamic tremor decoding sites were dorsolateral to optimal slowness decoding sites within individual patients

To verify the spatial relationship of optimal tremor and slowness decoding within patients, five additional right-handed patients (70.0 ± 8.9 years old; 2F, 3M; UPDRS III : 45.2 ± 9.5) underwent a modified version of the random-pattern task. Rather than acquire recordings from a stationary site within the STN, here we surveyed the entire length of the nucleus by systematically moving the electrodes between task trials in small, discrete steps (see *Materials and Methods*).

SVR models for tremor and slowness were then calculated as before, here incorporating recording data across all sites/trials within a single trajectory. Although decoding performance of neural models derived from these multi-site data exhibited a trend of lower performance than models trained on single-site recordings (Tremor: *r* = 0.196 ± 0.250 v. 0.094 ± 0.141; Slowness: *r* = 0.114 ± 0.193 v. 0.067 ± 0.144, Effective motor control: *r* = 0.138 ± 0.194 v. 0.055 ± 0.141), this difference was not found to be significant (*n* = 203 stationary microelectrode recording sites, *n* = 17 moving microelectrode recording sites, moving v. stationary data, LMM *β*s = −0.053 – - 0.102, *Z* = −0.766 – −1.057, *p* = 0.222 – 0.444). Of note, despite the lack of data at each recording site, whole-STN models demonstrated above-chance decoding performance for all three metrics (tremor: 8/17 recordings, slowness: 14/17 recordings, effective motor control: 10/17 recordings).

Recording sites along each trajectory were also reconstructed using imaging **(Fig. 4C)**, and site-specific metric decoding *r*-values were calculated by applying the whole-STN SVR model to individual site recordings (see *Materials and Methods, Neural Decoding of Behavioral Metrics)*.

Decoding performance (and the difference between tremor and slowness decoding performance) was then plotted across patients **(Fig. 4D)**. Across these additional MER recordings, tremor was optimally decoded at *x*=−13.0, *y*=−13.0, *z*=−5.0, while slowness was optimally decoded at *x*=−12.0, *y*=−13.0, *z*=−8.0. Again, optimal tremor decoding sites (AFNI’s “3dttest++”, *Z*=1.817, *p*=0.035) were dorsolateral to optimal slowness decoding sites (*Z*=2.050, *p*=0.020), with each metric peak occurring above chance. Taken together, tremor was found to be decoded dorsolaterally to slowness in both patient groups, using both across vs. within subject data.

### Cortical recordings also revealed distinct representations of tremor, slowness, and effective motor control

Of the 27 subjects who performed the fixed-pattern task, 10 subjects additionally had ECoG recordings from sensorimotor cortex. For this analysis, we considered only electrode contacts confirmed to be placed on primary motor or somatosensory cortices (motor: *n*=16 contacts, somatosensory: *n*=15, see *Materials and Methods*) based on intraoperative imaging. To determine if cortical recordings could be used to decode motor function and dysfunction on short timescales, SVR models were trained on ECoG signals recorded from sensorimotor cortex. There was no performance difference between ECoG decoding of tremor or slowness (*n* = 85 ECoG recordings across 27 session and 10 subjects, tremor: *r*=0.322±0.215, slowness: *r* =0.307±0.212, tremor v. slowness, LMM β = 0.015, *Z* = 0.557, *p*=0.578).

To understand which spectral features contributed to cortical decoding, SVR model weights were aggregated across all patients and recordings. Compared to metric-shuffled models, both tremor and slowness cortical decoding models had distinct distributions of weights across the spectrum **(Fig. 5A, B)**. When compared directly, cortical tremor and slowness models had opposing relationships in *α* / *β* (8–40 Hz, *p* < 0.027, permutation test), *γ*_*mid*_ (45–125, *p* < 0.023, permutation test), and *γ*_*high*_ (150–225 Hz, *p* < 0.015, permutation test) frequency bands **(Fig. 5C)**. Tremor models additionally had positive weights associated with *θ* frequencies (5–7 Hz, *p* = 0.003, permutation test). Altogether, although cortical signals supported equivalent decoding performance for tremor or slowness, decoding features were nonetheless distinct for each metric.

**Fig. 5.**
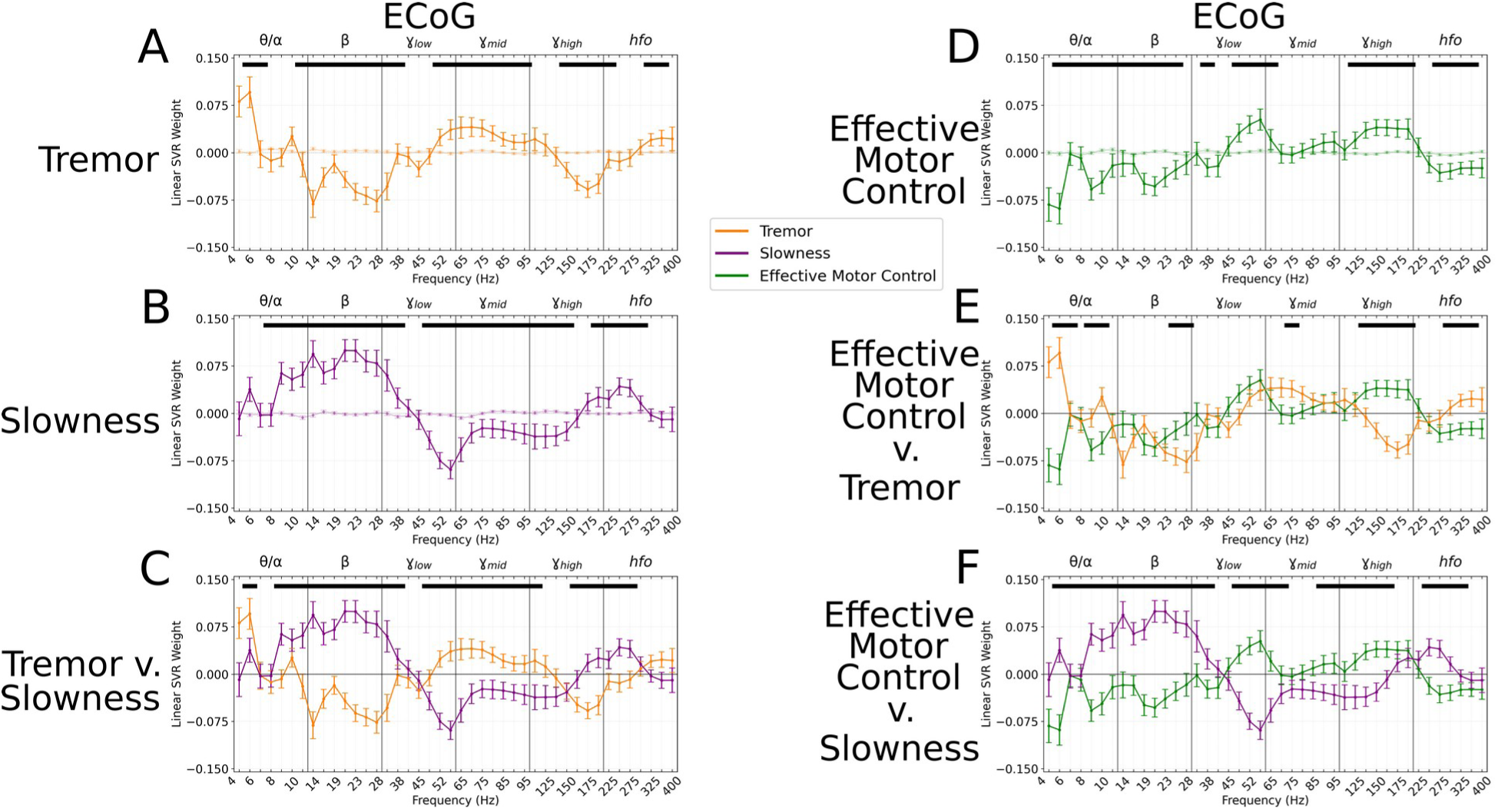
Cortical tremor and slowness decoding models exhibited opposing weights for multiple frequency bands. **(A)** Average cortical tremor decoding model coefficients for every recording along sensorimotor cortex. Colored solid lines indicate average weights, with positive/negative values reflecting a positive or negative relationship with tremor amplitude. Error bars indicate s.e.m. across subjects. Colored translucent lines represented mean weights (± s.e.m.) for SVR model weights calculated on shuffled metric data. Black solid lines (top) represented contiguous spectral features that were observed above chance. **(B)** Average slowness decoding model coefficients for every recording along sensorimotor cortex. **(C)** Average model coefficients for tremor and slowness. Black solid lines (top) represented contiguous spectral features that significantly differed between tremor and slowness decoding models. **(D)** Average cortical effective motor control decoding model coefficients for every recording along sensorimotor cortex. **(E)** Average model coefficients for effective motor control and tremor. **(F)** Average model coefficients for effective motor control and slowness.

On the other hand, effective motor control decoding performance (*r*=0.224 ± 0.177) was lower than tremor (effective motor control v. tremor, LMM β = −0.097, *Z* = −3.619, *p* = 2.96*10^−4^) and slowness decoding performance (effective motor control v. slowness, LMM β = −0.082, *Z* = - 3.062, *p* = 0.002). Nevertheless, effective motor control was represented in cortical decoding models by *γ*_*high*_ frequencies **(Fig. 5D)**. These *γ*_*high*_ features additionally appeared to differentiate effective motor control models from both tremor and slowness models (125–175 Hz, *p <* 0.026, permutation test) **(Fig. 5E, F)**. In addition, *α* / *β* (8–30 Hz, *p* < 0.001, permutation test) and *γ*_*low*_ (45–75 Hz, *p* < 0.010, permutation test) frequencies exhibited an opposing relationship between effective motor control and slowness, much like the interaction between tremor and slowness models **(Fig. 5F)**. Taken together, although at different *γ* frequencies, both STN (*γ*_*low*_) and sensorimotor cortex (*γ*_*high*_) exhibited features specific to effective motor control.

Finally, we analyzed whether motor features were selectively represented in different regions of cortex. ECoG recording sites (and their associated motor feature decoding performance) were plotted along a standard cortical surface **(Fig. 6A)**. Comparing tremor and slowness decoding performance by cortical anatomy revealed that slowness decoding had peaks in medial motor (*n* = 31 ECoG recording sites, *x*=−37.4, *y*=−15.8, *z*=+70.2; AFNI’s “3dttest++”, *T* = 2.332, *p* = 0.022) and somatosensory (*x*=−33.6, *y*=−37.5, *z*=+71.2; AFNI’s “3dttest++”, *T*=2.034, *p*=0.043) cortices. We also observed a trending tremor decoding peak in lateral somatosensory cortex (*x*=−46.4, *y*=−30.9, *z*=+67.1; AFNI’s “3dttest++”, *T*=1.650, *p*=0.10) **(Fig. 6B)**. Similar to the STN, effective motor control decoding performance (relative to tremor or slowness decoding) was found not to differ by cortical anatomy (AFNI’s “3dttest++”, *p*>0.05). In sum, within the context of the target-tracking task, optimal tremor cortical decoding sites were found along relatively more lateral somatosensory cortices, whereas optimal slowness decoding sites were observed to be relatively medial to tremor sites.

**Fig. 6.**
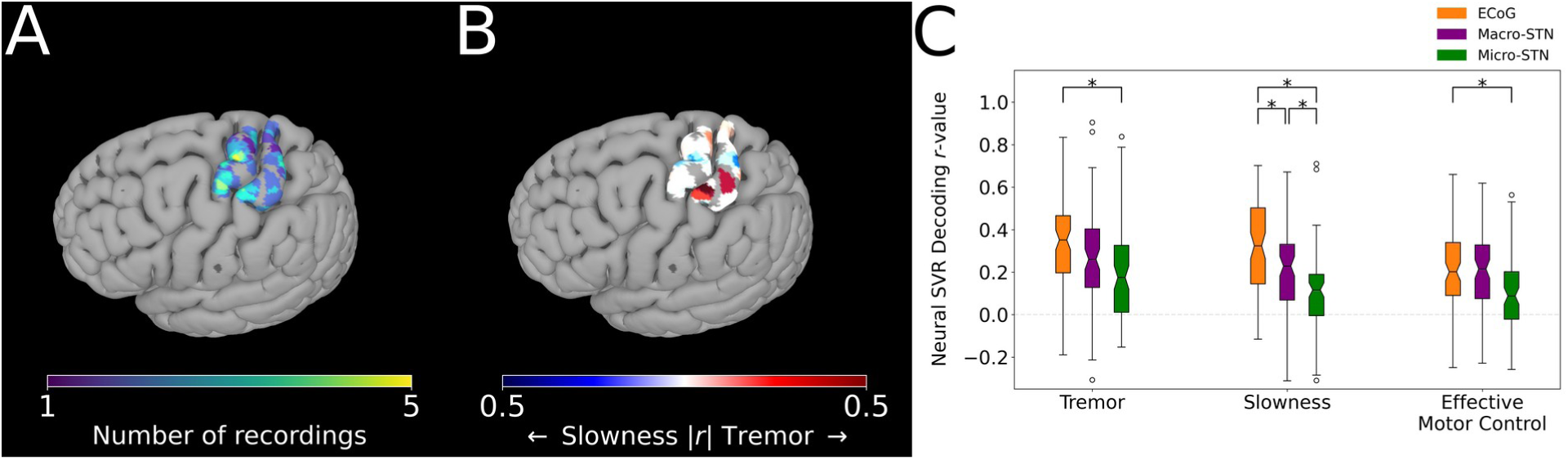
Cortical tremor and slowness decoding models were distributed throughout cortex, and generally were superior to STN decoding models. **(A)** Recording density of ECoG contacts on an MNI reference surface. **(B)** Difference in tremor vs. slowness decoding *r*-values for all cortical recordings. Warmer (red) colors indicate surface vertices where tremor decoding was superior, whereas cooler colors (blue) indicate where slowness decoding was superior. **(C)** Decoding performance across metrics and recording types. Box represents interquartile range (25th–75th percentile), while whiskers indicate 5th–95th percentile of data range. Brackets indicate significant (*, *p* < 0.05 in linear mixed model comparisons) differences in metric decoding *r*-values between recording types.

### Cortical decoding outperformed STN decoding

To understand whether motor (dys-)function was better represented in cortical signals, decoding performance was compared between patients with ECoG and micro-/macro-STN recordings (*n =* 10 subjects, 85 ECoG recordings, 81 microelectrode recordings, 81 macroelectrode recordings) **(Fig. 6C)**. ECoG tremor decoding models exhibited higher performance than micro-STN recordings (ECoG v. micro-STN, *r* = 0.322 ± 0.215 v. 0.204 ± 0.236, LMM β = 0.109, *Z* = 3.981, *p* = 6.86*10^−5^), but only trended higher relative to macro-STN recordings (ECoG v. macro-STN, *r* = 0.322 ± 0.215 v. 0.280 ± 0.219, LMM β = 0.034, *Z* = 1.232, *p* = 0.218). Slowness decoding performance on the other hand was higher in ECoG models relative to both macro-STN (ECoG v. macro-STN, *r* = 0.307 ± 0.212 v. 0.190 ± 0.205, LMM β = 0.114, *Z* = 4.382, *p* = 1.18*10^−5^) and micro-STN (ECoG v. micro-STN, *r* = 0.307 ± 0.212 v. 0.109 ± 0.175, LMM β = 0.195, *Z* = 7.511, *p* = 5.86*10^−14^) recordings. Like tremor, effective motor control ECoG decoding models exhibited superior decoding performance to micro-STN (ECoG v. micro-STN, *r* = 0.224 ± 0.177 v. 0.111 ± 0.176, LMM β = −0.114, *Z* = −4.409, *p* = 1.04*10^−5^), but not macro-STN (ECoG v. macro-STN, *r* = 0.224 ± 0.177 v. 0.206 ± 0.186, LMM β = 0.018, *Z* = 0.708, *p* = 0.479) recordings. In summary, recordings from sensorimotor cortex were superior to micro- and macro-STN recordings for decoding slowness, while cortical recordings were also superior to micro-STN recordings for decoding tremor and effective motor control. Overall, it appeared that cortical signals generally contained more relevant information for all three aspects of PD motor performance.

## DISCUSSION

In this study we quantified PD tremor and movement speed (slowness) in a structured motor task as surrogates for coarser clinical measurements of tremor and bradykinesia. We then decoded these metrics using linear models and mapped these results to basal ganglia and cortical anatomy. Our results demonstrate that tremor and bradykinesia were represented by different functional motifs with distinct localization in the STN and sensorimotor cortex. In addition, we contrasted these pathological states with periods of effective motor control, revealing unique markers of symptom-free states. To our knowledge this is the first study to not only characterize the behavioral interaction between tremor and slowness within a single behavioral context but also to compare directly each motor sign’s corresponding expression in neural activity, and further to compare this with relatively asymptomatic states. These results provide a holistic description of dynamic alternations in Parkinson’s symptoms which reveal specific neurophysiological biomarkers of non-pathological and distinct pathological states.

### The subthalamic nucleus exhibited clear functional and anatomical topography

We focused our neural decoding approach on two cardinal motor features of PD to isolate spectral features that reflected the expression of each motor sign/symptom. In the STN, tremor was characterized by lower frequency (*θ, α*) oscillations in microelectrodes, whereas slowness was characterized by the presence of *β* oscillations and the absence of broadband *γ* oscillations in macroelectrodes. Because *γ* frequency oscillations are commonly associated with hyperkinetic states *(19, 20)*, our slowness decoding results may be understood in part as an “anti-speed” neural model. Indeed, effective motor control was distinguished by *γ*_*low*_ frequency activity, highlighting the importance of *γ* frequency oscillations in effective (not simply dyskinetic) movements. These results fit within the broader STN literature, as some of these frequency bands in isolation (*θ*, *β*) have been found to correlate with clinical measures of Parkinson’s disease motor symptoms (tremor, bradykinesia) *(21–23)*. However, here we show directly the contrasting nature of distinct PD motor states both behaviorally and neurophysiologically, and highlight the dependence of these neurophysiological “fingerprints” on the particular neural recording technique.

We were also able to identify where tremor and slowness were more optimally decoded (*i*.*e*., where metric-specific spectral information was greatest). Within our STN microelectrode recordings, optimal tremor decoding sites were found to be located within dorsolateral STN whereas optimal slowness decoding sites were more centrally located within the STN. Considering the potential imprecision of pooling recording sites across multiple patients, optimal tremor decoding may have included activity from zona incerta (ZI), a stimulation site commonly thought to be critical for alleviating tremor *(24, 25)*. Indeed, optimal stimulation sites to alleviate tremor and bradykinesia may correspond to our dorsolateral-tremor/ventromedial-slowness topography *(11)*. While several groups have localized *β* frequency activity to dorsolateral STN, this has been observed to be located inferiorly to tremor-related higher frequency oscillations *(10, 26, 27)*. Our tremor/slowness-dorsal/ventral STN results correspond to prior work suggesting subdomains within the STN. In macaques, motor cortex projects to the dorsal portion of the STN, and the ventral STN receives projections from prefrontal cortex *(12)*. In humans, the ventral STN has been associated with stopping movement, while the dorsal STN is more associated with motor initiation and selection *(13, 28)*. Consequently, our tremor models fit within previous work finding tremor-frequency oscillations (*θ*) originating in the STN and propagating/synchronizing to motor cortex during tremor *(29, 30)*. Our anatomical results suggest that this propagation may be specific to dorsal STN. Slowness decoding models alternatively relied upon *β* activity in macroelectrode recordings, perhaps reflecting anti-kinetic *β* bursts relayed to ventral STN from inferior frontal or supplementary motor cortex *(31, 32)*. But while prior work did not directly compare the neuroanatomical substrate of distinct PD features within or across subjects, this work directly demonstrates how alternating motor features of PD manifest along these anatomical subdivisions.

### Cortical Parkinson’s disease motor decoding models were distinct from subthalamic models

In general, cortical recordings were equally capable of decoding tremor, slowness, and effective motor control, if. When comparing the feature weights of these decoding models, we observed generally opposing relationships in both *β* and *γ* frequency bands. As previous studies have shown that tremor decreases *β* oscillations across cortex *(33)*, and others have shown increased *γ* activity during hyperkinetic/dyskinetic states *(19)*, here we demonstrate a “push-pull” relationship between these frequency bands in the alternating expression of tremor and slowness, and when comparing slowness and effective motor control models. While cortical *β* frequency oscillations (and their desynchronization with movement) are well characterized in PD *(34)*, the functional role of *γ*_*high*_/*hfo* oscillations are less clear. Although these higher frequency oscillations overlap with phase-amplitude coupling peaks observed in cortex in unmedicated patients with PD *(35)*, our models for effective motor control states suggested that *γ*_*high*_ is specifically associated with more normal movement *(36)* and is not simply a marker for dyskinetic states *(19)*.

Although our cortical anatomical results (slowness-medial cortex/tremor-lateral cortex) did not reach significance, there is evidence for medial/midline cortex synchronizing with STN via *β* frequency oscillations *(37)*. The lateral aspect of our tremor results may also align with previous fMRI studies demonstrating lateral somatosensory and parietal cortical interactions with cerebellar thalamus during tremor *(38)*. Regardless, ECoG recordings across sensorimotor cortex provided relevant information for decoding tremor and slowness, as well as for identifying states of more effective motor function.

### Therapeutic Implications

With advances in technology, DBS aspires towards incorporating chronic neurophysiological recordings to help guide therapeutic stimulation *(39)*. While currently-applied closed-loop DBS paradigms trigger stimulation based on one or two frequency bands representing PD symptoms *(40)*, our results argue for the potential utility of a more fine-grained and targeted neurophysiological approach to PD state identification: more dorsal STN contacts may better sense signals reflecting tremor, while more ventral STN contacts may better identify signals corresponding to bradykinesia. Precise symptom-specific models could not only inform where to stimulate, but also when and *how* to stimulate (*i*.*e*., identifying stimulation settings to best treat tremor vs. bradykinesia). Most importantly, future neuromodulation paradigms could be derived not simply to disrupt pathological activity but actually to sustain the neurophysiological *γ* frequency “targets” of effective motor control. Looking ahead, chronic cortical recordings could work in concert with STN recordings to help identify precise motor states associated with specific aspects of disease expression.

### Limitations/Future directions

While our motor metrics correlated with UPDRS subscores, we recognize that our single intra-operative behavioral task does not capture all aspects of PD motor dysfunction. Further, while we focused on only spectral power measurements in each structure, future work may examine whether phase-based measures of synchrony across structures may potentially better decode motor states.

The accuracies of our multi-spectral models varied significantly across recordings and subjects, and the average decoding accuracies were somewhat low likely due to our inclusion of all recordings without pre-screening for subject PD type, precise recording location, signal quality, or signal information content. Nonetheless, decoding motor behavior was possible to some extent in all individuals, and was sufficiently successful to reveal significant differences between PD motor states. More accurate decoding models (arising from potentially different behavioral paradigms or analytic strategies) may further elucidate some of the PD state neural signatures observed here.

Given the nonuniform spatial sampling of imaging-based reconstructions, our imaging-based analyses may have lacked sufficient power to reveal smaller-scale or additional neurophysiological-anatomical relationships. The spatial resolution of recordings was also less precise in recordings with a large contact size (macroelectrodes, ECoG contacts). Moving forward, structural imaging techniques such as diffusion tensor imaging (DTI) across the STN and cortex could provide an anatomical anchor for the proposed circuits underlying the functional specificity of deep and cortical recordings. Nevertheless, with our parametric measurements of PD motor behavior of in the intraoperative setting, we were able to delineate the contrasting, push-pull relationship between neural states underlying tremor, bradykinesia, and effective motor control in both the STN and sensorimotor cortex.

## MATERIALS AND METHODS

### Study and Experimental Design

All patients undergoing routine, awake placement of deep brain stimulating electrodes for intractable, idiopathic Parkinson’s disease (PD) between June 2014 and December 2018 were invited to participate in this study. Patients were selected and offered DBS by a multi-disciplinary team based solely upon clinical criteria, and the choice of the target (STN vs. GPi) was made according to each patient’s particular circumstance (disease manifestations, cognitive status and goals) *(41)*. In this report, we focused on twenty-seven subjects (2F, 25M; aged 47.5– 78.5 years) undergoing STN DBS. Subjects were off all anti-Parkinsonian medications for at least 12 hours in advance of the surgical procedure (UPDRS Part III: 49.0 ± 13.3). Twelve subjects were considered tremor-dominant, and 13 subjects had average tremor UPDRS III scores 2 in their dominant hand *(42)*. Seventeen age-matched controls (14F, 3M; often patients’ partners; aged 48.5–79.2 years) also participated in this study (Patient v. control subjects ages, 65.2 ± 7.5 vs. 62.4 ± 10.0, Mann-Whitney U-test, *p*=0.319). Controls were required simply to be free of any diagnosed or suspected movement disorder and to have no physical limitation preventing them from seeing the display or manipulating the joystick. There was a strong male-bias in the patient population (2F, 25M) and a female preponderance in the control population (14F, 3M), reflecting weaker overall biases in the prevalence of PD and the clinical utilization of DBS therapy *(43–45)*. Both patients and control subjects were predominantly right-handed (patients: 25 right-handed, 2 left-handed; control subjects: 16 right-handed, 1 left-handed). Five additional right-handed patients aged 56.2–77.4 with PD (70.0 ± 8.9 years old; 2F, 3M; UPDRS III: 45.2 ± 9.5) underwent a modified version of tracking task where recordings were acquired throughout the entire STN. Of these five patients, two were tremor-dominant and only one had an average UPDRS III score > 2 in their dominant hand. Patients and control subjects agreeing to participate in this study signed informed consent, and experimental procedures were undertaken in accordance with an approved Rhode Island Hospital human research protocol (Lifespan IRB protocol #263157) and the Declaration of Helsinki.

### Surgical Procedure

Microelectrode recordings (MER) from the region of the STN of awake patients are routinely obtained in order to map the target area and guide DBS electrode implantation. A single bolus dose of short-acting sedative medication (typically propofol) was administered before the start of each procedure, at least 60–90 minutes prior to MER. The initial trajectory was determined on high-resolution (typically 3T) magnetic resonance images (MRI) co-registered with CT images demonstrating previously-implanted skull-anchor fiducial markers (version 3.0, FHC Inc., Bowdoin, ME, USA). Localization of the target relied upon a combination of direct and indirect targeting, utilizing the visualized STN as well as standard stereotactic coordinates relative to the anterior and posterior commissures. Appropriate trajectories to the target were then selected to avoid critical structures and to maximize the length of intersection with the STN. A 3-D printed stereotactic platform (STarFix micro-targeting system, FHC Inc.) was then created such that it could be affixed to these anchors, providing a precise trajectory to each target *(46)*. Microdrives (Alpha Omega Inc., Nazareth, Israel) were attached to the platform and then loaded with microelectrodes. Recordings were typically conducted along the anterior, center, and posterior trajectories (with respect to the initial MRI-determined trajectory) separated by 2 mm, corresponding to the axis of highest anatomical uncertainty based upon the limited visualization of the STN on MRI. For ten patients, electrocorticography (ECoG) strips were placed posteriorly along sensorimotor cortices through the same burr hole used for MER insertion to conduct intraoperative cortical recordings. MER began about 10–12 mm above the MRI-estimated target, which was chosen to lie near the inferior margin of the STN, about 2/3 of the distance laterally from its medial border. The STN was identified electrophysiologically as a hyperactive region typically first encountered about 3–6 mm above estimated target *(47)*. At variable intervals, when at least one electrode was judged to be within the STN, electrode movement was paused in order to assess neural activity and determine somatotopic correspondence, as per routine clinical practice. At these times, if patients were willing and able, additional recordings were obtained in conjunction with patient performance of the visual-motor task.

### High-Density STN survey

Five patients performed the visual-motor task in a modified paradigm. To start, clinical MER was conducted in typical fashion. Once the electrodes were judged to have exited the STN, the length of the STN recording span was calculated based on intraoperative neurophysiology. Based upon this length, the electrodes were automatically raised by the microdrives in pre-calculated steps in coordination with the visual-motor task, during the inter-trial-intervals. During this task, a separate control computer was used to coordinate the behavioral task with robotic control of the Alpha Omega neurophysiology and microdrive systems. Specifically, the FDA-approved C+ + Neuro Omega software development kit was compiled into a custom Python library that could communicate with the Neuro Omega systems with a ∼2 ms round-trip latency. From there, task-specific Python code enabled communication with the behavioral control system. To acquire microelectrode recordings that spanned the STN, the length of the STN was estimated based upon standard neurophysiological assessment (hyperactive region with frequent bursting activity and prominent low-frequency oscillations), and this length was divided by the number of task trials (typically 36). As the task was performed, the start of each inter-trial-interval was detected by the control computer, and every few trials (typically 3), a command was issued to raise the electrodes by the appropriate distance. The task re-commenced once drive movement was complete (typically ∼10 seconds later). This process continued until the subject completed the task and the microelectrodes had reached the top of the STN.

### Behavioral Task

We employed a visual-motor target tracking task with fixed- or random-patterns to estimate motor dysfunction in a quantitative and continuous fashion. Specifically, while subjects with PD reclined on the operating bed in a “lawn-chair” position, a joystick or stylus was positioned within their dominant hand, and a boom-mounted display was positioned within their direct line-of-sight at a distance of ∼1 meter. The fixed-pattern task was implemented in MonkeyLogic *(48–50)* and required subjects to follow a green target circle that moved smoothly around the screen by manipulating the joystick with the goal of keeping the white cursor within the circle **(Fig. 1A)**. The target circle followed one of several possible paths (invisible to the subject), with each trial lasting 10–30 seconds. Each session consisted of up to 36 trials (∼13 minutes of tracking data), and subjects performed 1–4 sessions during the operation. To address potential confounds in the fixed-pattern task such as implicitly learning patterns, a second version of the task was later created and implemented in NIMH MonkeyLogic *(51)*. Instead of fixed-pattern paths, this version implemented randomly generated patterns unique to each trial with paths optimized to maintain a near-steady target movement speed *(16)*. While the goal remained the same (following the onscreen target with a patient-controlled cursor), the cursor was controlled here by the patient moving a stylus with their dominant hand along a tablet. For either task version, control subjects performed this task in an extra-operative setting.

### Behavioral Metric Calculation

Regardless of the task version performed, tremor amplitude and movement speed were calculated from the cursor x- and y-traces. Tremor amplitude was calculated from 3–10 Hz band-pass filtered x- and y-cursor traces, specifically as the Euclidean mean of the envelopes of each cursor dimension’s analytic signal (*TM*_*x*_(*t*)=∥*C*_*ax*_∥). Movement speed was calculated from cursor traces low-pass filtered at 3 Hz to remove the influence of tremor, and quantified as the Euclidean change of cursor position over time 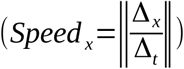. Both metrics were averaged into 7 second non-overlapping epochs (*n* = 6498 epochs across 44 subjects) to maintain consistency with our previous decoding approach *(17)*. To standardize movement speed within subjects, movement speed epochs within a session were min-max normalized into a measure of “slowness,” where 0=highest speed and 1=lowest speed.

Finally, to quantify effective motor control in our task, effective motor control was quantified as the inverse of tremor and slowness measures, relative to the entire session. Specifically, tremor and slowness were additionally min-max transformed in an inverted fashion, where 0=highest tremor/lowest speed and 1=lowest tremor/highest speed. Each epoch’s “effective motor control” measure was then calculated as 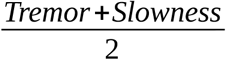, where performance values of 0 indicated the most symptomatic states (tremor, slowness) whereas values of 1 indicated optimal motor performance.

### Behavioral Metric Analysis

Behavioral metrics (tremor and slowness) were compared across control and PD populations using a linear mixed model to account for each subject’s asymmetric contribution of epochs: *y*_*metric*_=*X* _*population*_*β*+*Zu*+*ϵ*, where *y*_*metric*_ represented each epoch’s metric amplitude, *X*_*population*_ represented categorical labels of populations and associated fixed-effect regression coefficients (*β*), *Z* represented the subject-specific random intercepts and their associated random effect coefficients (*u*), and *ϵ* represented the residuals. To understand the interactions and optimal timescales of tremor and slowness, each metric was calculated within smaller, 100 ms epochs. Linear mixed models were used to calculate the correlation between tremor and slowness across the entirety of each subject’s behavioral data using the following model: *y*_*tremor*_=*X*_*slowness*_ *β*+*Zu*+*ϵ*, where *y*_*tremor*_ represented each epoch’s tremor amplitude, *X*_*slowness*_ represented the epoch’s simultaneous measurement of slowness and the fixed-effect regression coefficient (*β*), and *Zu* represented the same subject-specific random-intercept design as above.

To determine the timescales of each metric’s expression, autocorrelograms were calculated similarly across each PD subject’s behavioral data. The average full-width half-maximum (FWHM) of the autocorrelograms were subsequently identified as the minimum time necessary to label motor metric data as a “symptomatic” period. Tremor or slowness were considered “symptomatic” if they were above the 95th percentile of aggregate control data, and sustained symptomatic periods were defined as those persisting beyond the population metric FWHM continuously. For effective motor control, epochs were labeled “symptomatic” if they were above the median of the PD subject’s session distribution distribution, and persisting beyond the PD population FWHM continuously.

### Neurophysiological Signals and Analysis

Microelectrode signals were recorded using “NeuroProbe” tungsten electrodes (Alpha Omega), and macroelectrode signals were recorded from circumferential contacts 3 mm above the microelectrode tips. ECoG signals were acquired using Ad-Tech 8-contact subdural strips with 10 mm contact-to-contact spacing (Ad-Tech Medical, Racine, WI). All signals were acquired at 22–44 kHz and synchronized using Neuro Omega data acquisition systems (Alpha Omega). Microelectrode impedances were typically 400–700 k*Ω*, macroelectrode impedances were typically 2–20 k*Ω*, and ECoG contact impedances were typically 10–30 k*Ω*. Patients performed up to 4 sessions of the task, with microelectrodes positioned at different depths for each session. As microelectrodes were not independently positionable (and macroelectrodes were 3 mm above microelectrode tips along the trajectory), some signals may have necessarily been acquired outside of the STN (see Fig. 4). All signals were nevertheless considered and analyzed.

Neural data were analyzed using the “numpy/scipy” Python 3 environment (https://numpy.org/, https://www.scipy.org/) *(52, 53)*. Offline, ECoG signals were re-referenced to a common median reference within a strip *(54)*. All resulting signals were bandpass filtered between 2–600 Hz, and notch filtered at 60 Hz and its harmonics. Timeseries were then Z-scored and artifacts above 4 standard deviations (often representing movement artifact) were removed. These resulting timeseries were then downsampled to 1 kHz. Timeseries were bandpass filtered using a Morlet wavelet convolution (wave number 7) at 1 Hz intervals, covering 3–400 Hz. The instantaneous power and phase at each frequency was then determined by the Hilbert transform. To analyze broad frequency bands, we grouped frequencies into canonical ranges: *θ* /*α*: 3–12 Hz, *β*_*low*_: 12–20 Hz, *β*_*high*_: 20–30 Hz, *γ*_*low*_: 30–60 Hz, *γ*_*mid*_: 60–100 Hz, *γ*_*high*_: 100–200 Hz, and *hfo* (high frequency oscillations): 200–400 Hz.

### Imaging-based reconstruction of recording sites

Patients underwent pre-, intra- and post-operative imaging per routine clinical care. Preoperatively, stereotactic protocol magnetic resonance (MR) images were obtained (Siemens Vario 3.0 T scanner) that included T1- and T2-weighted sequences (T1: MPRAGE sequence; TR: 2530 ms, TE: 2.85 ms, matrix size: 512 × 512, voxels: 0.5 × 0.5 mm^2^ in-plane resolution, 224 sagittal slices, 1 mm slice thickness; T2: SPACE sequence, TR: 3200 ms, TE: 409 ms, matrix size: 512 × 512, voxels: 0.5 × 0.5 mm^2^ in-plane resolution, 224 sagittal slices, 1 mm slice thickness). Pre-, intra-, and post-operative (in some cases) computed tomography (CT) scans were also acquired (Extra-Op CT: GE Lightspeed VCT Scanner; Tube voltage: 120 kV, Tube current: 186 mA, data acquisition diameter: 320 mm, reconstruction diameter: 250 mm, matrix size: 512 × 512 voxels, 0.488 × 0.488 mm^2^ in-plane resolution, 267 axial slices, 0.625 mm slice thickness; Intra-Op CT: Mobius Airo scanner, Tube voltage: 120 kV, Tube current: 240 mA, data acquisition diameter: 1331 mm, reconstruction diameter: 337 mm, matrix size: 512 × 512 voxels, 0.658 × 0.658 mm^2^ in-plane resolution, 182 axial slices, 1 mm slice thickness).

Postoperative MR images (Siemens Aera 1.5 T scanner, T1: MPRAGE sequence, TR: 2300 ms, TE: 4.3 ms, matrix size: 256 × 256 voxels, 1.0 × 1.0 mm^2^ in-plane resolution, 183 axial slices, 1 mm slice thickness, specific absorption rate < 0.1 W/g) were typically obtained 1–2 days after the operation to confirm proper final electrode location. To reconstruct recording locations, MR and CT images were co-registered using the FHC Waypoint Planner software. The raw DICOM images and the linear transform matrices were exported and applied to reconstructed image volumes using the AFNI command “3dAllineate,” bringing them into a common coordinate space *(55, 56)*. Microelectrode depths were calculated by combining intraoperative recording depth information with electrode reconstructions obtained from postoperative images using methods described previously *(57, 58)*. To determine the anatomical distribution of microelectrode recording sites across patients, preoperative T1-weighted MR images were registered to a T1-weighted MNI reference volume (MNI152 T1 2009c) using the AFNI command “3dQwarp” *(59)*. The resulting patient-specific transformation was then applied to recording site coordinates. MNI-warped recording coordinates were then assessed for proximity to structures such as the STN as delineated on the MNI PD25 atlas *(60–62)*. ECoG contacts were segmented from intraoperative CT volumes, and were then projected onto individual cortical surface reconstructions generated from preoperative T1 volumes *(63–66)*. Individual cortical surface reconstructions were co-registered to a standard Desikan-Destrieux surface parcellation *(67–69)*. Contacts were labeled and grouped as “premotor cortex,” “motor cortex,” “somatosensory cortex,” or “parietal cortex” if they contained the following anatomical parcellation labels:

- Premotor cortex/PMC : ctx_lh_G_front_sup, ctx_lh_G_front_middle
- Motor cortex/MC : ctx_lh_G_precentral
- Somatosensory cortex/SC : ctx_lh_G_postcentral
- Posterior Parietal cortex/PPC : ctx_lh_G_parietal_sup, ctx_lh_G_pariet_inf-Supramar

If a contact had more than one label (8/80 contacts), they were removed from further analysis. For this analysis, only contacts within sensorimotor cortex (motor or somatosensory cortex) were considered.

### Neural decoding of behavioral metrics

To investigate whether STN or cortical activity could be used to estimate co-occuring behavioral metrics, support vector regression (SVR) with “scikit-learn” (“linear” kernel with default parameters: *C* = 1.0, *tol* = 0.001; neural inputs and behavioral outputs min-max scaled from 0–1) was applied towards multi-spectral decoding of tremor or slowness *(70)*. In order to use neural features across the entire spectrum for neural decoding, spectral power estimates for each canonical band (*θ* /*α, β*_*low*_, *β*_*high*_, *γ*_*low*_, *γ*_*mid*_, *γ*_*high*_, *hfo*) were further subdivided into 7 sub-bands for a total of 42 spectral features across 3–400 Hz *(17)*. SVR models trained on a single electrode’s spectral features were fit using 100-fold Monte Carlo cross-validation using a 2:1 train/test split of temporal epochs within a task session. Model performance was assessed by linear regression (specifically, the Pearson *r*-value) between the empirical/observed and predicted metric distributions. To verify that these decoded results were not spurious, a separate set of SVR models were fit using the same parameters as above but with a shuffled correspondence between behavioral metric data and neurophysiological signals in the training set.

When assessing whether one type of metric (e.g. tremor) was preferentially decoded within a single type of recording (e.g. microelectrodes), SVR *r*-values were compared using the following linear mixed model: *y*_*r*−*value*_=*X*_*metric*_*β*+*Zu*+*ϵ*, where *y*_*r-value*_ represented SVR decoding *r*-values from a single recording and metric, *X*_*metric*_ represented categorical labels of metrics and associated fixed-effect regression coefficients (*β*), *Z* represented the subject-specific random intercepts and their associated random effect coefficients (*u*), and *ϵ* represented the residuals. When investigating whether one type of recording was superior at decoding a single metric, *r*-values were compared using the following linear mixed model: *y*_*r*_=*X*_*recording*_*β*+*Zu*+*ϵ*, where *y*_*r*_ represented SVR decoding *r*-values from a single recording and metric, *X*_*recording*_ represented categorical labels of recording types and associated fixed-effect regression coefficients (*β*), and *Zu* represented the same subject-specific random intercepts model as above.

When comparing cross-metric model performance (*i*.*e*. determining the ability of a model trained on tremor to decode slowness), performance was assessed by linear regression between the model’s predicted metric (tremor) distribution and the co-occurring alternate metric (slowness) distribution. To compare the relative performance of tremor-trained models on decoding slowness, *r*-values were compared within recording type using the following linear mixed model: *y*_*r*_=*X*_*metric*_*β*+*Zu*+*ϵ*, where *y*_*r*_ represented the SVR decoding *r*-value, *X*_*metric*_ represented the categorical labels of either the model’s trained metric (tremor) or the alternate metric (slowness) and their associated fixed-effect regression coefficients (*β*), *Z* represented the subject-specific random intercepts and their associated random effect coefficients (*u*), and *ϵ* represented the residuals.

Because these SVR models used a linear kernel, we were able to extract SVR model coefficients (“weights”) to understand which spectral features were used to decode behavioral metrics. As linear SVR estimates of behavioral metrics (*Y*_*behavior*_) are a combination of neural weights (*W*_*neural*_) and power estimate (*X*_*neural*_) inputs (*Y*_*behavior*_=*W*_*neural*_·*X*_*neural*_+*intercept*), positive weights described the association between the presence of a specific frequency band with higher metric output values. Conversely, negative weights described the absence of a neural feature when metric output values were high.

To test whether specific clusters of features (>= 3 contiguous spectral features) were consistently weighted across recordings, the distribution of each feature’s SVR model weights (averaged over 3 adjacent features) across recordings were compared to the distribution of metric-shuffled SVR model weights using a contiguity-sensitive permutation test *(17)*. Over 10000 iterations, each recording’s SVR weight values were shuffled across the two models (empirical vs. shuffled), and the difference between individual feature distributions across electrodes was assessed using a paired *t*-test. The empirical *T* -statistic for each feature distribution (*e*.*g*., microelectrode *β*_1−3_ empirical v. shuffled) was then compared to the null *T* -statistic distribution. The probability of the observed difference in *T*-statistics was calculated empirically from the null *T*-statistic distribution. This procedure was also used for understanding whether specific features were weighted differently for specific metric decoding models (*e*.*g*., microelectrode *β*_1−3_ tremor v. slowness).

For datasets collected using the within-subject, high-density STN survey, SVR models were trained using recordings throughout the STN. Specifically, recordings at each depth were split into 2:1 train:test sets and aggregated for whole-STN SVR model fitting. *r*-values from SVR models trained on high-density microelectrode data were compared to *r-*values from stationary microelectrode data using the following linear mixed model: *y*_*r*_=*X*_*experiment*_*β*+*Zu*+*ϵ*, where *y*_*r*_ represented the SVR decoding *r*-value, *X*_*experiment*_ represented the categorical labels of experiment type and their associated fixed-effect regression coefficients (*β*), *Z* represented the subject-specific random intercepts and their associated random effect coefficients (*u*), and *ϵ* represented the residuals.

To determine if whole-STN models were able to decode metrics above chance, *r-*value distributions from empiric and metric-shuffled decoding models were compared using the Wilcoxon test. To understand if specific recording sites contained information specific to individual metrics, metrics were additionally estimated at each depth by applying the whole-STN SVR model weights to spectral features recorded at each depth. From there, *r*-values for each recording/depth were calculated between estimated and observed metrics.

### Anatomical analysis of metric decoding

To compare whether specific motor features were better decoded in different regions of the STN, tremor and slowness *r*-values were plotted in MNI coordinate space. All voxels and their associated *r*-values with MER recordings were then compared with a voxel-wise paired *t*-test with AFNI’s “3dttest++” function. Each resulting voxel had an associated *Z*-statistic that was generated from 10000 permutations of shuffling tremor and dataset *r*-values across voxels using AFNI’s Equitable Thresholding and Clustering (ETAC) algorithm *(71)*. Briefly, ETAC was used to estimate dataset-specific empirical statistical (e.g. *T*-statistic) and cluster-size (number of adjacent voxels) thresholds for significant results by running several (10000) permutations of testing with shuffled data. However, due to the relatively low number of voxels per recording dataset (1 mm^3^ per recording site compared to whole-brain coverage typically acquired with fMRI scans), no distinct clusters were isolated by the ETAC algorithm. Nevertheless, we still reported the voxelwise *Z*-scores (and the corresponding *p*-values from the Gaussian probability density function) from the permutation. For cortical surface-based comparisons between tremor and slowness *r*-values, “3dttest++” was also used, although without the ETAC algorithm as it is not currently implemented for surface-based datasets. Thus instead of *Z*-scores computed by ETAC, we examined the *T* -statistic from “3dttest++” and its associated *p*-value. To correct for cluster size, we considered only clusters that were above the cluster size corresponding to *p*=0.05 in “slow_surf_clustsim.py”, where 1000 simulations of clustering were performed on all cortical surface vertices containing ECoG contacts. Similarly to STN results, we were unable to identify any significant clusters at *p*=0.05 due to spatial undersampling of the cortical surface but still reported the vertexwise *T* statistics. Colormaps for cortical figures were obtained from https://github.com/snastase/suma-colormaps.

### Statistical Analysis

Data in text are represented as mean ± standard deviation. All statistical tests, unless otherwise specified, were carried out in the “scipy” python environment. *P*-values were adjusted for multiple comparisons wherever appropriate using the Benjamini-Hochberg procedure with *q*=0.05 *(72)*. When comparing data aggregated across multiple subjects, linear mixed models were performed using the “statsmodels” python toolbox to disentangle the effect of interest (continuous or categorical) from the random effects/unequal contributions of each subject’s dataset *(73, 74)*. All linear mixed models were random intercepts models, where each random intercept corresponded to a subject’s dataset, and generally followed the formula of *y*=*Xβ*+*Zu*+*ϵ*, where *y* represented the outcome variable, *X* represented the continuous or categorical predictor variables, *β* represented the fixed-effect regression coefficients, *Z* represented the subject-specific random intercepts, *u* represented the random-effects regression coefficients, and *ϵ* represented the residuals of the model fit. Once a model was fit, fixed-effect *p*-values were calculated from *Z*-scored parameter estimates (fixed-effect coefficients divided by their standard errors) against the normal distribution.

## Acknowledgments

We are grateful for the generous participation of our patients in this study. We thank Kelsea Laubenstein-Parker for technical assistance, Karina Bertsch for administrative support, and Ann Duggan-Winkle for clinical support. We also thank James Yu, Minkyu Ahn, David Segar, Tina Sankhla, and Daniel Shiebler for helping develop the motor task experiment. Finally, we thank Menem Andrea and Alpha Omega for adding computer-controlled microdrive capabilities.

## Funding

National Institutes of Health Training Grant NINDS T32MH020068 (PML)

Doris Duke Clinical Scientist Development Award #2014101 (WFA)

National Institutes of Health COBRE Award: NIGMS P20 GM103645 (PI: Jerome Sanes) supporting WFA

Neurosurgery Research and Education Foundation grant (WFA)

Lifespan Norman Prince Neurosciences Institute

Brown University Robert J. and Nancy D. Carney Institute for Brain Science

Part of this research was conducted using computational resources and services at the Center for Computation and Visualization at Brown University, with funding provided by an NIH Office of the Director grant S10OD025181

WFA has received proprietary equipment and technical support for unrelated research through the Medtronic external research program.

## Author contributions

Conceptualization: PML, SL, WFA

Methodology: PML, SL, DEA, DDL

Investigation: PML, SL, DEA, UA, WFA

Formal Analysis: PML

Funding acquisition: WFA

Supervision: SL, UA, WFA

Writing – original draft: PML

Writing – review & editing: PML, SL, DEA, DDL, UA, WFA

## Competing interests

The authors have patents and patent applications broadly relevant to Parkinson’s disease (but not directly based upon this work). WFA has received proprietary equipment and technical support for unrelated research through the Medtronic external research program.

## Data and materials availability

The datasets supporting the current study have not been deposited in a public repository because they contain patient information but are available along with analysis code from the corresponding authors upon request.

## References and Notes

1. J. Parkinson, An essay on the shaking palsy. 1817. J Neuropsychiatry Clin Neurosci 14, 223–236; discussion 222 (2002).

2. M. J. Armstrong, M. S. Okun, Diagnosis and Treatment of Parkinson Disease: A Review. JAMA 323, 548–560 (2020).

3. W. C. Koller, Pharmacologic Treatment of Parkinsonian Tremor. Arch Neurol 43, 126–127 (1986).

4. K. Sethi, Levodopa unresponsive symptoms in Parkinson disease. Movement Disorders 23, S521–S533 (2008).

5. P. Limousin, P. Krack, P. Pollak, A. Benazzouz, C. Ardouin, D. Hoffmann, A. L. Benabid, Electrical stimulation of the subthalamic nucleus in advanced Parkinson’s disease. N. Engl. J. Med. 339, 1105–1111 (1998).

6. M. Katz, M. S. Luciano, K. Carlson, P. Luo, W. J. Marks, P. S. Larson, P. A. Starr, K. A. Follett, F. M. Weaver, M. B. Stern, D. J. Reda, J. L. Ostrem, CSP 468 study group, Differential effects of deep brain stimulation target on motor subtypes in Parkinson’s disease. Ann. Neurol. 77, 710–719 (2015).

7. S. Little, M. Beudel, L. Zrinzo, T. Foltynie, P. Limousin, M. Hariz, S. Neal, B. Cheeran, H. Cagnan, J. Gratwicke, T. Z. Aziz, A. Pogosyan, P. Brown, Bilateral adaptive deep brain stimulation is effective in Parkinson’s disease. J. Neurol. Neurosurg. Psychiatr. 87, 717–721 (2016).

8. S. Little, E. Tripoliti, M. Beudel, A. Pogosyan, H. Cagnan, D. Herz, S. Bestmann, T. Aziz, B. Cheeran, L. Zrinzo, M. Hariz, J. Hyam, P. Limousin, T. Foltynie, P. Brown, Adaptive deep brain stimulation for Parkinson’s disease demonstrates reduced speech side effects compared to conventional stimulation in the acute setting. J. Neurol. Neurosurg. Psychiatry 87, 1388–1389 (2016).

9. D. Piña-Fuentes, J. M. C. van Dijk, J. C. van Zijl, H. R. Moes, T. van Laar, D. L. M. Oterdoom, S. Little, P. Brown, M. Beudel, Acute effects of Adaptive Deep Brain Stimulation in Parkinson’s disease. Brain Stimulation (2020), doi:10.1016/j.brs.2020.07.016.

10. I. Telkes, A. Viswanathan, J. Jimenez-Shahed, A. Abosch, M. Ozturk, A. Gupte, J. Jankovic, N. F. Ince, Local field potentials of subthalamic nucleus contain electrophysiological footprints of motor subtypes of Parkinson’s disease. Proc. Natl. Acad. Sci. U.S.A. 115, E8567–E8576 (2018).

11. H. Akram, S. N. Sotiropoulos, S. Jbabdi, D. Georgiev, P. Mahlknecht, J. Hyam, T. Foltynie, P. Limousin, E. De Vita, M. Jahanshahi, M. Hariz, J. Ashburner, T. Behrens, L. Zrinzo, Subthalamic deep brain stimulation sweet spots and hyperdirect cortical connectivity in Parkinson’s disease. NeuroImage 158, 332–345 (2017).

12. W. I. A. Haynes, S. N. Haber, The Organization of Prefrontal-Subthalamic Inputs in Primates Provides an Anatomical Substrate for Both Functional Specificity and Integration: Implications for Basal Ganglia Models and Deep Brain Stimulation. J. Neurosci. 33, 4804–4814 (2013).

13. C. P. Mosher, A. N. Mamelak, M. Malekmohammadi, N. Pouratian, U. Rutishauser, Distinct roles of dorsal and ventral subthalamic neurons in action selection and cancellation. Neuron 109, 869–881.e6 (2021).

14. W.-J. Neumann, H. Schroll, A. L. de Almeida Marcelino, A. Horn, S. Ewert, F. Irmen, P. Krause, G.-H. Schneider, F. Hamker, A. A. Kühn, Functional segregation of basal ganglia pathways in Parkinson’s disease. Brain 141, 2655–2669 (2018).

15. W. S. Gibson, A. E. Rusheen, Y. Oh, M.-H. In, K. R. Gorny, J. P. Felmlee, B. T. Klassen, S. J. Jung, H.-K. Min, K. H. Lee, H. J. Jo, Symptom-specific differential motor network modulation by deep brain stimulation in Parkinson’s disease. Journal of Neurosurgery 1, 1–9 (2021).

16. J. B. Sanderson, J. H. Yu, D. D. Liu, D. Amaya, P. M. Lauro, A. D’Abreu, U. Akbar, S. Lee, W. F. Asaad, Multi-Dimensional, Short-Timescale Quantification of Parkinson’s Disease and Essential Tremor Motor Dysfunction. Front. Neurol. 11 (2020), doi:10.3389/fneur.2020.00886.

17. M. Ahn, S. Lee, P. Lauro, E. Schaeffer, U. Akbar, W. Asaad, Rapid motor fluctuations reveal short-timescale neurophysiological biomarkers of Parkinson’s disease. J. Neural Eng. (2020), doi:10.1088/1741-2552/abaca3.

18. P. Mazzoni, A. Hristova, J. W. Krakauer, Why Don’t We Move Faster? Parkinson’s Disease, Movement Vigor, and Implicit Motivation. J. Neurosci. 27, 7105–7116 (2007).

19. N. C. Swann, C. de Hemptinne, S. Miocinovic, S. Qasim, S. S. Wang, N. Ziman, J. L. Ostrem, M. San Luciano, N. B. Galifianakis, P. A. Starr, Gamma Oscillations in the Hyperkinetic State Detected with Chronic Human Brain Recordings in Parkinson’s Disease. J. Neurosci. 36, 6445–6458 (2016).

20. R. Lofredi, W.-J. Neumann, A. Bock, A. Horn, J. Huebl, S. Siegert, G.-H. Schneider, J. K. Krauss, A. A. Kühn, E. Vaadia, Ed. Dopamine-dependent scaling of subthalamic gamma bursts with movement velocity in patients with Parkinson’s disease. eLife 7, e31895 (2018).

21. Y. Nie, H. Luo, X. Li, X. Geng, A. L. Green, T. Z. Aziz, S. Wang, Subthalamic dynamic neural states correlate with motor symptoms in Parkinson’s Disease. Clinical Neurophysiology (2021), doi:10.1016/j.clinph.2021.07.022.

22. N. Asch, Y. Herschman, R. Maoz, C. Aurbach-Asch, D. Valsky, M. Abu-Snineh, D. Arkadir, E. Linetsky, R. Eitan, O. Marmor, H. Bergman, Z. Israel, Independently together: subthalamic theta and beta opposite roles in predicting Parkinson’s tremor. Brain Commun (2020), doi:10.1093/braincomms/fcaa074.

23. W.-J. Neumann, K. Degen, G.-H. Schneider, C. Brücke, J. Huebl, P. Brown, A. A. Kühn, Subthalamic synchronized oscillatory activity correlates with motor impairment in patients with Parkinson’s disease. Movement Disorders 31, 1748–1751 (2016).

24. C. Reck, E. Florin, L. Wojtecki, H. Krause, S. Groiss, J. Voges, M. Maarouf, V. Sturm, A. Schnitzler, L. Timmermann, Characterisation of tremor-associated local field potentials in the subthalamic nucleus in Parkinson’s disease. European Journal of Neuroscience 29, 599–612 (2009).

25. P. Plaha, S. Khan, S. S. Gill, Bilateral stimulation of the caudal zona incerta nucleus for tremor control. Journal of Neurology, Neurosurgery & Psychiatry 79, 504–513 (2008).

26. I. Tamir, D. Wang, W. Chen, J. L. Ostrem, P. A. Starr, C. de Hemptinne, Eight cylindrical contact lead recordings in the subthalamic region localize beta oscillations source to the dorsal STN. Neurobiology of Disease 146, 105090 (2020).

27. B. C. M. van Wijk, A. Pogosyan, M. I. Hariz, H. Akram, T. Foltynie, P. Limousin, A. Horn, S. Ewert, P. Brown, V. Litvak, Localization of beta and high-frequency oscillations within the subthalamic nucleus region. Neuroimage Clin 16, 175–183 (2017).

28. W. Chen, C. de Hemptinne, A. M. Miller, M. Leibbrand, S. J. Little, D. A. Lim, P. S. Larson, P. A. Starr, Prefrontal-Subthalamic Hyperdirect Pathway Modulates Movement Inhibition in Humans. Neuron (2020), doi:10.1016/j.neuron.2020.02.012.

29. J. Hirschmann, C. J. Hartmann, M. Butz, N. Hoogenboom, T. E. Özkurt, S. Elben, J. Vesper, L. Wojtecki, A. Schnitzler, A direct relationship between oscillatory subthalamic nucleus–cortex coupling and rest tremor in Parkinson’s disease. Brain 136, 3659–3670 (2013).

30. P. M. Lauro, S. Lee, U. Akbar, W. F. Asaad, Subthalamic-Cortical Network Reorganization during Parkinson’s Tremor. J. Neurosci. (2021), doi:10.1523/JNEUROSCI.0854-21.2021.

31. R. Hannah, V. Muralidharan, K. K. Sundby, A. R. Aron, Temporally-precise disruption of prefrontal cortex informed by the timing of beta bursts impairs human action-stopping. NeuroImage 222, 117222 (2020).

32. A. Oswal, C. Cao, C.-H. Yeh, W.-J. Neumann, J. Gratwicke, H. Akram, A. Horn, D. Li, S. Zhan, C. Zhang, Q. Wang, L. Zrinzo, T. Foltynie, P. Limousin, R. Bogacz, B. Sun, M. Husain, P. Brown, V. Litvak, Neural signatures of hyperdirect pathway activity in Parkinson’s disease. Nat Commun 12, 5185 (2021).

33. S. E. Qasim, C. de Hemptinne, N. C. Swann, S. Miocinovic, J. L. Ostrem, P. A. Starr, Electrocorticography reveals beta desynchronization in the basal ganglia-cortical loop during rest tremor in Parkinson’s disease. Neurobiology of Disease 86, 177–186 (2016).

34. N. C. Rowland, C. de Hemptinne, N. C. Swann, S. Qasim, S. Miocinovic, J. Ostrem, R. T. Knight, P. A. Starr, Task-related activity in sensorimotor cortex in Parkinson’s disease and essential tremor: changes in beta and gamma bands. Front. Hum. Neurosci. 9 (2015), doi:10.3389/fnhum.2015.00512.

35. C. de Hemptinne, E. S. Ryapolova-Webb, E. L. Air, P. A. Garcia, K. J. Miller, J. G. Ojemann, J. L. Ostrem, N. B. Galifianakis, P. A. Starr, Exaggerated phase-amplitude coupling in the primary motor cortex in Parkinson disease. Proc. Natl. Acad. Sci. U.S.A. 110, 4780–4785 (2013).

36. P. Fischer, A. Pogosyan, D. M. Herz, B. Cheeran, A. L. Green, J. Fitzgerald, T. Z. Aziz, J. Hyam, S. Little, T. Foltynie, P. Limousin, L. Zrinzo, P. Brown, H. Tan, E. Vaadia, Ed. Subthalamic nucleus gamma activity increases not only during movement but also during movement inhibition. eLife 6, e23947 (2017).

37. A. Sharott, A. Gulberti, S. Zittel, A. A. T. Jones, U. Fickel, A. Münchau, J. A. Köppen, C. Gerloff, M. Westphal, C. Buhmann, W. Hamel, A. K. Engel, C. K. E. Moll, Activity Parameters of Subthalamic Nucleus Neurons Selectively Predict Motor Symptom Severity in Parkinson’s Disease. J. Neurosci. 34, 6273–6285 (2014).

38. M. F. Dirkx, H. Zach, A. van Nuland, B. R. Bloem, I. Toni, R. C. Helmich, Cerebral differences between dopamine-resistant and dopamine-responsive Parkinson’s tremor. Brain 142, 3144–3157 (2019).

39. R. Gilron, S. Little, R. Perrone, R. Wilt, C. de Hemptinne, M. S. Yaroshinsky, C. A. Racine, S. S. Wang, J. L. Ostrem, P. S. Larson, D. D. Wang, N. B. Galifianakis, I. O. Bledsoe, M. San Luciano, H. E. Dawes, G. A. Worrell, V. Kremen, D. A. Borton, T. Denison, P. A. Starr, Long-term wireless streaming of neural recordings for circuit discovery and adaptive stimulation in individuals with Parkinson’s disease. Nature Biotechnology, 1–8 (2021).

40. Y. M. Kehnemouyi, K. B. Wilkins, C. M. Anidi, R. W. Anderson, M. F. Afzal, H. M. Bronte-Stewart, Modulation of beta bursts in subthalamic sensorimotor circuits predicts improvement in bradykinesia. Brain 144, 473–486 (2021).

41. U. Akbar, W. F. Asaad, A Comprehensive Approach to Deep Brain Stimulation for Movement Disorders. R I Med J (2013) 100, 30–33 (2017).

42. J. Jankovic, M. McDermott, J. Carter, S. Gauthier, C. Goetz, L. Golbe, S. Huber, W. Koller, C. Olanow, I. Shoulson, Variable expression of Parkinson’s disease: a base-line analysis of the DATATOP cohort. The Parkinson Study Group. Neurology 40, 1529–1534 (1990).

43. E. Accolla, E. Caputo, F. Cogiamanian, F. Tamma, S. Mrakic-Sposta, S. Marceglia, M. Egidi, P. Rampini, M. Locatelli, A. Priori, Gender differences in patients with Parkinson’s disease treated with subthalamic deep brain stimulation. Movement Disorders 22, 1150–1156 (2007).

44. G.-M. Hariz, T. Nakajima, P. Limousin, T. Foltynie, L. Zrinzo, M. Jahanshahi, K. Hamberg, Gender distribution of patients with Parkinson’s disease treated with subthalamic deep brain stimulation; a review of the 2000–2009 literature. Parkinsonism & Related Disorders 17, 146–149 (2011).

45. K. Rumalla, K. A. Smith, K. A. Follett, J. M. Nazzaro, P. M. Arnold, Rates, causes, risk factors, and outcomes of readmission following deep brain stimulation for movement disorders: Analysis of the U.S. Nationwide Readmissions Database. Clin Neurol Neurosurg 171, 129–134 (2018).

46. P. E. Konrad, J. S. Neimat, H. Yu, C. C. Kao, M. S. Remple, P.-F. D’Haese, B. M. Dawant, Customized, miniature rapid-prototype stereotactic frames for use in deep brain stimulator surgery: initial clinical methodology and experience from 263 patients from 2002 to 2008. Stereotact Funct Neurosurg 89, 34–41 (2011).

47. R. E. Gross, P. Krack, M. C. Rodriguez-Oroz, A. R. Rezai, A.-L. Benabid, Electrophysiological mapping for the implantation of deep brain stimulators for Parkinson’s disease and tremor. Mov. Disord. 21 Suppl 14, S259–283 (2006).

48. W. F. Asaad, E. N. Eskandar, Achieving behavioral control with millisecond resolution in a high-level programming environment. J. Neurosci. Methods 173, 235–240 (2008).

49. W. F. Asaad, E. N. Eskandar, A flexible software tool for temporally-precise behavioral control in Matlab. J. Neurosci. Methods 174, 245–258 (2008).

50. W. F. Asaad, N. Santhanam, S. McClellan, D. J. Freedman, High-performance execution of psychophysical tasks with complex visual stimuli in MATLAB. J Neurophysiol 109, 249–260 (2013).

51. J. Hwang, A. R. Mitz, E. A. Murray, NIMH MonkeyLogic: Behavioral control and data acquisition in MATLAB. J. Neurosci. Methods 323, 13–21 (2019).

52. C. R. Harris, K. J. Millman, S. J. van der Walt, R. Gommers, P. Virtanen, D. Cournapeau, E. Wieser, J. Taylor, S. Berg, N. J. Smith, R. Kern, M. Picus, S. Hoyer, M. H. van Kerkwijk, M. Brett, A. Haldane, J. F. del Río, M. Wiebe, P. Peterson, P. Gérard-Marchant, K. Sheppard, T. Reddy, W. Weckesser, H. Abbasi, C. Gohlke, T. E. Oliphant, Array programming with NumPy. Nature 585, 357–362 (2020).

53. P. Virtanen, R. Gommers, T. E. Oliphant, M. Haberland, T. Reddy, D. Cournapeau, E. Burovski, P. Peterson, W. Weckesser, J. Bright, S. J. van der Walt, M. Brett, J. Wilson, K. J. Millman, N. Mayorov, A. R. J. Nelson, E. Jones, R. Kern, E. Larson, C. J. Carey, İ. Polat, Y. Feng, E. W. Moore, J. VanderPlas, D. Laxalde, J. Perktold, R. Cimrman, I. Henriksen, E. A. Quintero, C. R. Harris, A. M. Archibald, A. H. Ribeiro, F. Pedregosa, P. van Mulbregt, SciPy 1.0: fundamental algorithms for scientific computing in Python. Nat Methods, 1–12 (2020).

54. Y. Liu, W. G. Coon, A. de Pesters, P. Brunner, G. Schalk, The effects of spatial filtering and artifacts on electrocorticographic signals. J. Neural Eng. 12, 056008 (2015).

55. R. W. Cox, AFNI: Software for Analysis and Visualization of Functional Magnetic Resonance Neuroimages. Computers and Biomedical Research 29, 162–173 (1996).

56. X. Li, P. S. Morgan, J. Ashburner, J. Smith, C. Rorden, The first step for neuroimaging data analysis: DICOM to NIfTI conversion. Journal of Neuroscience Methods 264, 47–56 (2016).

57. P. M. Lauro, N. Vanegas-Arroyave, L. Huang, P. A. Taylor, K. A. Zaghloul, C. Lungu, Z. S. Saad, S. G. Horovitz, DBSproc: An open source process for DBS electrode localization and tractographic analysis. Hum. Brain Mapp. (2015), doi:10.1002/hbm.23039.

58. P. M. Lauro, S. Lee, M. Ahn, A. Barborica, W. F. Asaad, DBStar: An Open-Source Tool Kit for Imaging Analysis with Patient-Customized Deep Brain Stimulation Platforms. SFN 96, 13–21 (2018).

59. V. Fonov, A. Evans, R. McKinstry, C. Almli, D. Collins, Unbiased nonlinear average age-appropriate brain templates from birth to adulthood. NeuroImage 47, S102 (2009).

60. Y. Xiao, S. Beriault, G. B. Pike, D. L. Collins, Multicontrast multiecho FLASH MRI for targeting the subthalamic nucleus. Magn Reson Imaging 30, 627–640 (2012).

61. Y. Xiao, V. Fonov, S. Bériault, F. Al Subaie, M. M. Chakravarty, A. F. Sadikot, G. B. Pike, D. L. Collins, Multicontrast unbiased MRI atlas of a Parkinson’s disease population. Int J Comput Assist Radiol Surg 10, 329–341 (2015).

62. Y. Xiao, V. Fonov, M. M. Chakravarty, S. Beriault, F. Al Subaie, A. Sadikot, G. B. Pike, G. Bertrand, D. L. Collins, A dataset of multi-contrast population-averaged brain MRI atlases of a Parkinson’s disease cohort. Data Brief 12, 370–379 (2017).

63. A. M. Dale, B. Fischl, M. I. Sereno, Cortical Surface-Based Analysis: I. Segmentation and Surface Reconstruction. NeuroImage 9, 179–194 (1999).

64. B. Fischl, D. H. Salat, E. Busa, M. Albert, M. Dieterich, C. Haselgrove, A. van der Kouwe, R. Killiany, D. Kennedy, S. Klaveness, A. Montillo, N. Makris, B. Rosen, A. M. Dale, Whole Brain Segmentation: Automated Labeling of Neuroanatomical Structures in the Human Brain. Neuron 33, 341–355 (2002).

65. Z. S. Saad, R. C. Reynolds, SUMA. Neuroimage 62, 768–773 (2012).

66. M. S. Trotta, J. Cocjin, E. Whitehead, S. Damera, J. H. Wittig, Z. S. Saad, S. K. Inati, K. A. Zaghloul, Surface based electrode localization and standardized regions of interest for intracranial EEG. Human Brain Mapping 39, 709–721 (2018).

67. B. D. Argall, Z. S. Saad, M. S. Beauchamp, Simplified intersubject averaging on the cortical surface using SUMA. Human Brain Mapping 27, 14–27 (2006).

68. R. S. Desikan, F. Ségonne, B. Fischl, B. T. Quinn, B. C. Dickerson, D. Blacker, R. L. Buckner, A. M. Dale, R. P. Maguire, B. T. Hyman, M. S. Albert, R. J. Killiany, An automated labeling system for subdividing the human cerebral cortex on MRI scans into gyral based regions of interest. NeuroImage 31, 968–980 (2006).

69. C. Destrieux, B. Fischl, A. Dale, E. Halgren, Automatic parcellation of human cortical gyri and sulci using standard anatomical nomenclature. Neuroimage 53, 1–15 (2010).

70. F. Pedregosa, G. Varoquaux, A. Gramfort, V. Michel, B. Thirion, O. Grisel, M. Blondel, P. Prettenhofer, R. Weiss, V. Dubourg, J. Vanderplas, A. Passos, D. Cournapeau, M. Brucher, M. Perrot, É. Duchesnay, Scikit-learn: Machine Learning in Python. Journal of Machine Learning Research 12, 2825–2830 (2011).

71. R. W. Cox, Equitable Thresholding and Clustering: A Novel Method for Functional Magnetic Resonance Imaging Clustering in AFNI. Brain Connectivity 9, 529–538 (2019).

72. Y. Benjamini, Y. Hochberg, Controlling the False Discovery Rate: A Practical and Powerful Approach to Multiple Testing. Journal of the Royal Statistical Society. Series B (Methodological) 57, 289–300 (1995).

73. M. J. Lindstrom, D. M. Bates, Newton-Raphson and EM Algorithms for Linear Mixed-Effects Models for Repeated-Measures Data. Journal of the American Statistical Association 83, 1014–1022 (1988).

74. S. Seabold, J. Perktold, statsmodels: Econometric and statistical modeling with python. 9th Python in Science Conference (2010).

